# A brain-tumour axis links chronic stress to gastric cancer progression through NPY signalling

**DOI:** 10.64898/2026.06.28.735136

**Authors:** Xiaomin Huang, Zhonghao Wu, Qi Wang, Cheng Wei, Jingyu Wang, Xinjie Ning, Rao Fu, Linxiang Lan, Changhua Zhang, Yulong He, Siqiang Ren, Brian G Oliver, Hui Chen, Alexei Verkhratsky, Jianqin Niu, Chenju Yi

**Author notes:** Correspondence to: Jianqin Niu: No.30 Gaotanyan Street, Shapingba District, Chongqing 400038, China;., Chenju Yi: No. 628 Zhenyuan Road, Guangming District, Shenzhen 518107, China;. The authors contributed equally (Xiaomin Huang, Zhonghao Wu).

## Abstract

Chronic stress promotes gastric tumour growth, but the central neural pathways remain largely enigmatic. Here, we identify a brain-tumour axis, implicating the central amygdala (CeA) activity in promoting gastric cancer growth through sympathetic neuropeptide Y (NPY) signalling occurring within the tumour. Using clinical datasets, patient samples, pathway tracing, electrophysiology and chemogenetic manipulation, we demonstrate that stress selectively increases NPY Y1 receptor (NPY1R) synthesis in the patient’s gastric cancer tissue; moreover, the NPY1R density correlates with advanced cancer stage and poor prognosis. In mice, daily constraint chronic stress enhances CeA excitability and promotes sympathetic neurotransmitter norepinephrine release within the tumour microenvironment. Norepinephrine, in turn, increases tumour endogenous NPY/NPY1R production and the activation of downstream MAPK/Erk1/2 signalling pathway, which drives cancer cell proliferation and invasion. Manipulating CeA neuronal activity alone can regulate tumour growth, whereas selectively blocking tumour NPY1R prevents stress-induced tumour progression *in vivo* and cancer cell migration *in vitro*. This study identifies a central-to-peripheral circuit linking chronic stress to gastric cancer growth, and highlights NPY1R as a potential therapeutic target.

## Introduction

Gastric cancer remains the fifth most common malignancy and the fifth leading cause of cancer-related mortality worldwide, largely owing to its marked heterogeneity and early metastatic potential ^1^. Despite substantial advances in the research on genomics and tumour immunology, clinical outcomes remain suboptimal, particularly due to recurrence and drug resistance ^1,2^. In this context, emerging evidence has redefined tumours as dynamic entities subjected to regulation by the nervous system, rather than isolated pathologies within cancer cells, which gave rise to a new field, cancer neuroscience ^3–6^. Neural fibres infiltrate solid tumours and provide inputs through the local release of neurotransmitters, neuromodulators and neurotrophic factors which modify tumour behaviour ^5,7^, including tumour growth, nutrient metabolism, immune evasion, and metastatic expansion ^8–10^. However, the majority of studies have focused on local nerve-tumour interactions at the tumour site ^7^, whereas the role of central nervous system in regulation of tumour progression through defined neural efferent pathways remains largely unknown.

The stomach represents a unique model for studying brain-tumour interactions due to its dense and well-defined bidirectional autonomic and sensory innervation. Physiologically, gastric function is under central control of tightly coordinated sympathetic and parasympathetic (vagal) pathways, which provide a direct anatomical and functional interface through which brain activity influences local tissue homeostasis and organ function ^11–13^. This balance is disrupted in disease and during tumourigenesis. Vagal signalling promotes gastric cancer development, acting through muscarinic receptor 3 ^14^, whereas sympathetic signalling contributes to tumour vascularisation and malignant progression utilising adrenergic pathways ^17,18^. Consequently, activation of the sympathetic nervous system alone is associated with tumour exacerbation across multiple cancer models ^18,19^. However, it is unclear which brain regions link stress and subsequently regulate efferent sympathetic outputs that remodel the tumour microenvironment in stomach cancer.

Chronic stress, commonly manifested by anxiety and depression, is highly prevalent among cancer patients and is associated with increased cancer incidence and progression. It also accelerates cancer progression and promotes treatment resistance ^20–23^. Stress responses are typically mediated through the activation of the hypothalamic-pituitary-adrenal (HPA) axis and the sympathetic-adreno-medullary system, resulting in increased circulating glucocorticoids and catecholamines ^24,25^, which enhance gastric tumour growth and promotes metastasis in the mouse model of chronic restraint stress ^26^. However, hormonal changes in the blood alone do not adequately explain the spatial specificity and magnitude of tumour progression. Beyond fast-acting neurotransmitters, neuropeptides produced locally at the tumour site may provide a sustained mechanistic bridge between central chronic stress responses and long-term tumour remodelling. Here, we focus on neuropeptide Y (NPY), a conserved 36-amino acid peptide widely expressed in the nervous system; NPY regulates stress responses when produced in the brain, while in the periphery, it controls autonomic function and cell proliferation ^27–32^. The central amygdala (CeA) is a key hub for processing negative emotions, including stress ^33,34^, by integrating cortical and thalamic inputs to regulate efferent autonomic output ^35^. In the amygdala, NPY is known to regulate anxiety-related behaviours ^36^, specifically through NPY Y1 receptors (NPYR1) ^27^. In the tumour tissue, locally produced NPY is involved in tumour progression in several cancer types ^37^; although NPY mechanisms were not characterised in gastric cancer, especially under chronic stress conditions.

Therefore, we hypothesised that chronic stress engages the CeA-associated efferent sympathetic activation, which in turn activates NPY/NPY1R signalling to promote gastric cancer growth. We aimed to define a brain-tumour axis to provide mechanistic insight into how chronic stress drives gastric cancer progression. To address this, we combined *in vivo* retrograde trans-synaptic tracing using pseudorabies virus and chemogenetic manipulation of CeA neuronal activity, and *in vitro* assays, to map the anatomical connectivity between stress-responsive CeA and gastric cancers and identify how sympathetic neurotransmitter norepinephrine (NE) regulates endogenous tumour NPY signalling.

## Results

### 1. Elevated NPY1R expression and increased sympathetic nerve infiltration are closely associated with gastric cancer progression in human patients

To confirm whether the NPY signalling pathway is involved in gastric cancer progression, we first performed bioinformatics analysis of publicly available databases. Using transcriptomic data of patients with stomach adenocarcinoma (STAD) in The Cancer Genome Atlas (TCGA) database, we evaluated the association between gastric cancer and NPY, as well as NPY major receptor subtypes (NPY1R, NPY2R, NPY4R, NPY4R2, NPY5R, and NPY6R). Both NPY and NPY1R were significantly elevated in gastric cancer tissues (Fig. 1A). Further analysis across different TNM stages showed that increased NPY1R expression, but not but not NPY itself or the other receptor subtypes, significantly correlates with the advancement of TNM stages (Fig. 1B, *p* = 0.037) and with the decreases survival rate of gastric cancer patients (Fig. 1C; Extended Data Fig.1A-L). Therefore, NPY1R is closely associated with gastric cancer progression and prognosis.

**Fig. 1.**
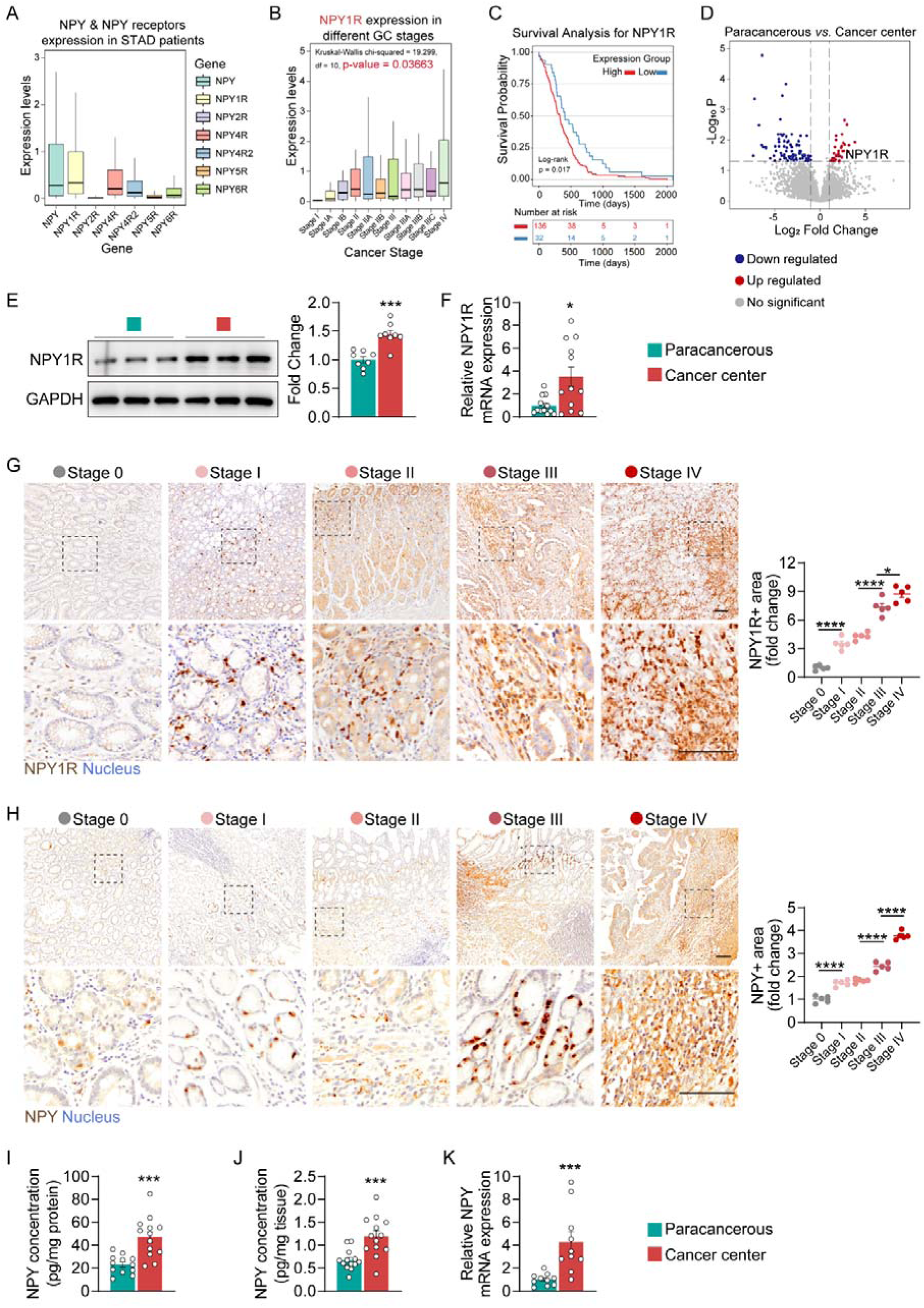
Increased NPY1R level closely associated with gastric cancer progression in patients. **(A)** Gene expression profiles of NPY and its receptor subtypes (NPY1R, NPY2R, NPY4R, NPY4R2, NPY5R, and NPY6R) in the stomach adenocarcinoma cohort in the Cancer Genome Atlas. (B) The expression of NPY1R across different TNM stages in the stomach adenocarcinoma cohort in the Cancer Genome Atlas. (C) Association between NPY1R expression and overall survival in patients with gastric cancer. (D) Increased NPY1R expression in gastric cancer tissues from patients by RNA-seq analysis. (E) NPY1R protein level (n = 8) and (F) NPY1R mRNA expression (n = 12) in gastric cancer and paracancerous tissues. (G) Representative images and quantification of NPY1R stain in gastric cancer tissues from TNM stages 0-IV, scale bar: 100 μm, n = 5 patients. (H) Representative images and quantification of NPY staining in gastric cancer tissues from TNM stages 0-IV, scale bar: 100 μm, n = 5 patients. (I) NPY protein level normalised to total protein in gastric cancer and paracancerous tissues from gastric cancer patients, n = 13 patients. (J) ELISA analysis of NPY protein level in paracancerous and tumour tissues from gastric cancer patients, normalised to tissue weight. n = 13 patients. (K) NPY mRNA expression in cancer and paracancerous tissues, n = 10 patients. Data are presented as mean ± SEM, and were analysed by two-tailed *t* tests (2 groups) or one-way ANOVA with Tukey post hoc tests (> 2 groups), \**p* < 0.05, \*\**p* < 0.01, \*\*\**p* < 0.001, \*\*\*\**p* < 0.0001.

To validate the above bioinformatics analysis results, RNA-seq analysis was performed for cancer tissues and paracancerous non-malignant tissues from gastric cancer The NPY1R levels in gastric cancer tissues were significantly higher than those in paracancerous tissues (Fig. 1D). Both Western blot and qPCR showed higher levels of NPY1R mRNA and protein expression in tumour tissues compared to paracancerous tissues (Fig. 1E,F, protein *p* < 0.001, mRNA *p* < 0.05). Next, we performed immunohistochemical analysis and showed that NPY1R expression in gastric cancer tissues progressively increased with advancing TNM stage (Fig. 1G), with significant increases observed from stage 0 through stage IV. These results further confirmed that NPY1R level is closely associated with gastric cancer progression.

Then, we examined NPY distribution using immunohistochemical staining. The number of NPY-positive cells in tumour tissues increased with advancing TNM stages (Fig. 1H). ELISA confirmed higher NPY protein levels in tumour tissues than those in paracancerous tissues when normalised to total protein (Fig. 1I, *p* < 0.001) or by tissue weight (Fig. 1J, *p* < 0.001). NPY mRNA expression was also upregulated in tumour tissues compared with paracancerous tissues (Fig. 1K, *p* < 0.001). However, plasma NPY concentrations in gastric cancer patients were similar to those of healthy controls (Extended Data Fig.1M,N). The above results suggest that the elevated NPY in gastric cancer tissues is primarily derived from the local tumour microenvironment rather than from the circulation.

To evaluate the dynamic changes of neural components during cancer progression, we measured nerve markers in gastric cancer tissues of different TNM stages, including the neuronal marker microtubule-associated protein 2 (MAP2), the sympathetic nerve marker tyrosine hydroxylase (TH), and the vagal nerve marker choline acetyltransferase (ChAT). The levels of MAP2 and TH in gastric cancer tissues progressively increased with advancing TNM stages (Fig. 2A,B). In contrast, the vagal nerve marker ChAT in gastric cancer tissues did not change across different TNM stages (Fig. 2C). These results suggest that the neural and sympathetic nerve fibre infiltration is associated with gastric cancer progression.

**Fig. 2.**
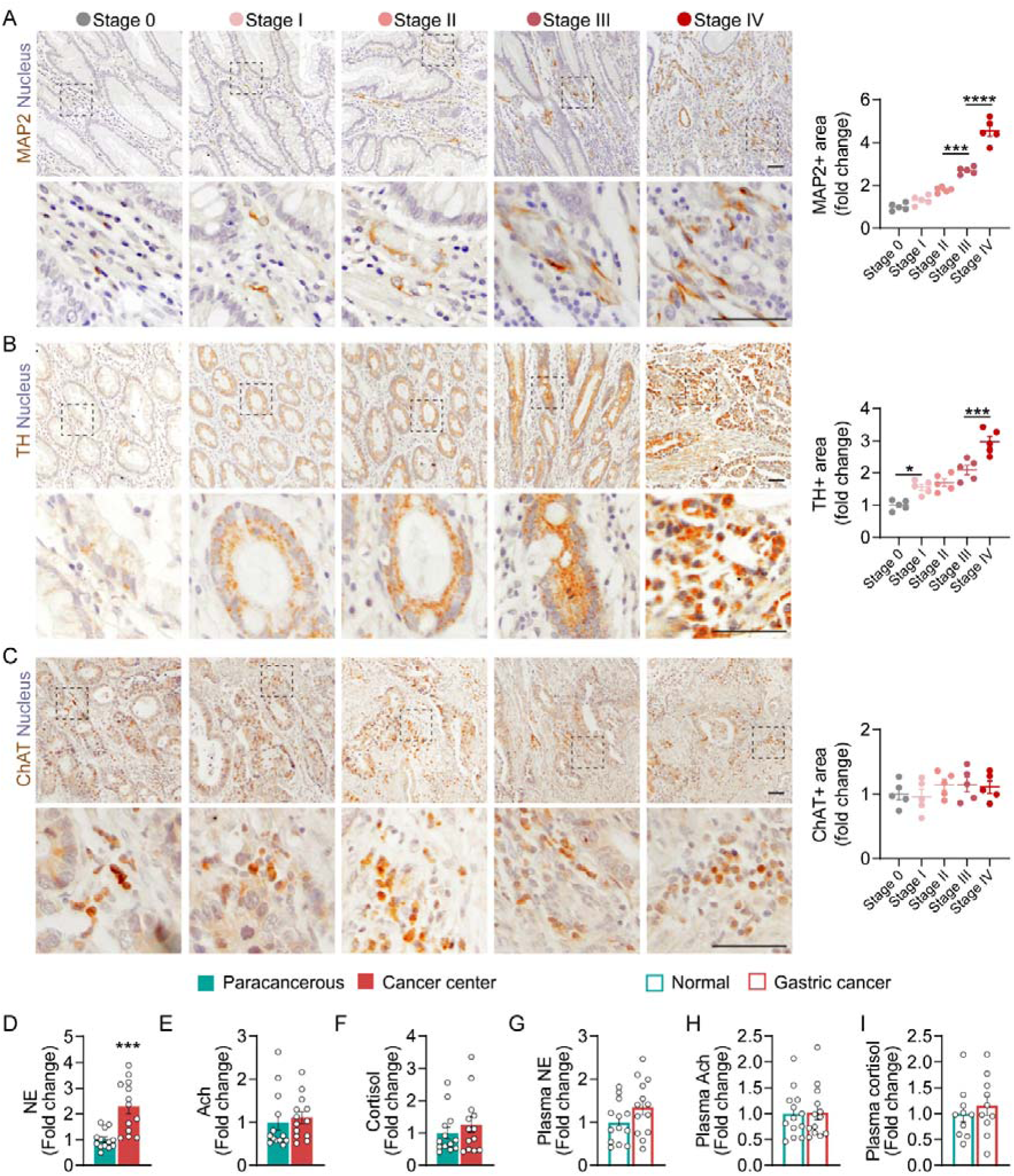
Increased sympathetic nerve infiltration and increased release of sympathetic neurotransmitter norepinephrine in gastric cancer tissue from patients. (A) Representative images of microtubule associated protein 2 (MAP2) staining in gastric cancer of TNM stages 0-IV, scale bar: 100 μm. Right: Quantification of MAP2+ area in gastric cancer of TNM stages 0-IV. n = 5. (B) Representative images of tyrosine hydroxylase (TH) staining in gastric cancer from TNM stages 0-IV, scale bar: 100 μm. Right: TH+ area in gastric cancer tissues from TNM stages 0-IV. n = 5. (C) Representative images of choline acetyl transferase (ChAT) staining in gastric cancer of TNM stages 0-IV, scale bar: 100 μm. Right: ChAT+ area in gastric cancer from TNM stages 0-IV. N = 5. (D) NE levels in paracancerous and gastric cancer tissues from patients, n = 13. (E) Ach levels in paracancerous and gastric cancer tissues from patients, n = 13. (F) Cortisol level in paracancerous and gastric cancer tissues from patients, n = 13. Plasma (G) NE, (H) Ach and (I) cortisol levels in gastric cancer patients and healthy controls, n = 13. Data are presented as mean ± SEM, and were analysed by two-tailed *t* tests (2 groups) or one-way ANOVA with Tukey post hoc tests (> 2 groups), \**p* < 0.05, \*\**p* < 0.01, \*\*\**p* < 0.001, \*\*\*\**p* < 0.0001.

Finally, we measured the changes in the sympathetic neurotransmitter norepinephrine (NE), the vagal neurotransmitter acetylcholine (ACh), and the stress hormone cortisol in the cancer, paracancerous tissues, and plasma. NE levels in cancer tissue were significantly higher than those in paracancerous tissues (Fig. 2D, *p* < 0.001). However, no significant differences in ACh or cortisol were observed between cancer and paracancerous tissues (Fig. 2E,F). Moreover, plasma levels of NE, ACh, and cortisol did not differ between patients with gastric cancer and healthy controls (Fig. 2G-I). These results suggest that only the sympathetic neurotransmitter is significantly elevated in the local microenvironment of gastric cancer tissues, consistent with the increased sympathetic nerve infiltration.

### 2. Chronic restraint stress increased NPY1R expression and promoted gastric cancer progression in mice

Brain NPY level can be regulated by stress ^32,38,39^, but less is known about its response in gastric cancer. Thus, we investigated how chronic stress affects the NPY/NPY1R signalling in gastric cancer and relates to disease progression. We used a nude mouse model of chronic restraint stress combined with inguinal subcutaneous xenotransplantation of HGC27 human gastric cancer cells (Fig. 3A). On day 7 after stress initiation, HGC27 cells were subcutaneously injected into the inguinal region of BALB/c-nude mice. At the same time, NPY1R-specific inhibitor BIBO3304 (5 mg/kg, intraperitoneal (i.p.) injection once every 2 days) was initiated until the endpoint (day 35). Three groups were established: Ctrl (tumour+vehicle), Stress (tumour+chronic restraint stress+vehicle), and Stress+BIBO (tumour+chronic restraint stress+BIBO3304 treatment). Since BIBO3304 does not cross the blood-brain barrier ^40^, central NPY effects are excluded.

**Fig. 3.**
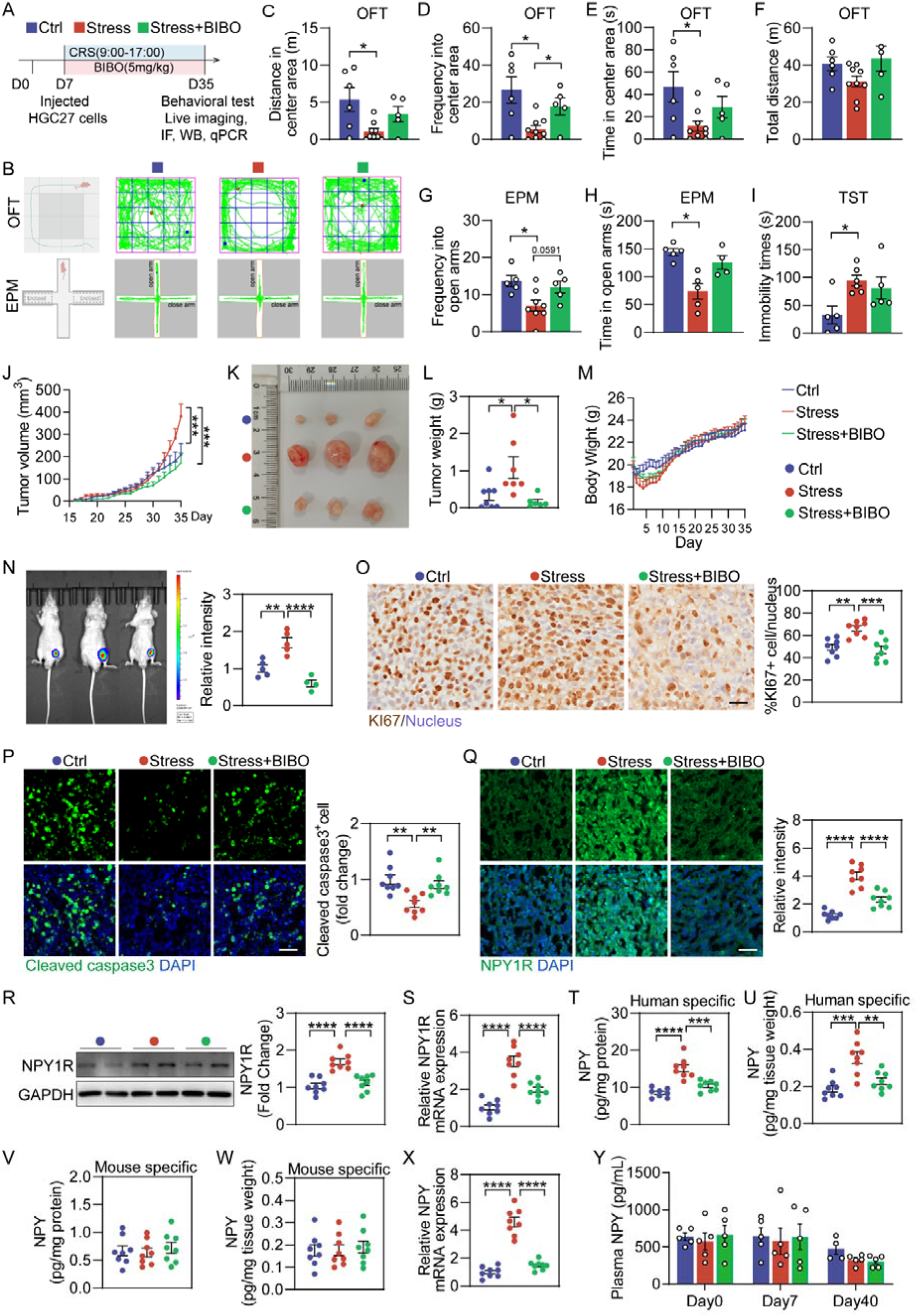
Chronic restraint stress enhances tumour growth and activates the NPY/NPY1R axis in a mouse model of gastric cancer. (A) Schematic diagram of the modelling of chronic restraint stress in tumour-bearing mice. (B) Representative travel trace of the open field test (OFT) and elevated plus maze (EPM). (C) Distance travelled, (D) number of entries, and (E) the time spent in the centre area, and (F) total distance in the OFT, n = 4-9. (G) Number of entries and (H) time spent in the open arms in the EPM. n = 5-8. (I) Immobility time in the tail suspension test (TST), n = 5-6. (J) Changes in tumour volume over time, n= 8-13. (K) Representative images of tumours at the endpoint. (L) Tumour weight, n = 6-8. (M) Body weight changes. n = 17-18. (N) Representative images of *in vivo* bioluminescence imaging of HGC27-luciferase labelled tumours. Right: bioluminescence intensity. n = 4-5. (O) Representative images of KI67 staining in the tumours, scale bar: 20 μm. Right: the percentage of KI67-positive cells relative to the number of cell nuclei, n = 8. (P) Representative images of cleaved caspase3 staining in tumours, scale bar: 50 μm. Right: cleaved caspase3-positive cell number, n = 8. (Q) Representative images of NPY1R immunostaining in tumours, scale bar: 50 μm. Right: NPY1R intensity. n = 8. (R) NPY1R protein levels in tumours, n = 8. (S) NPY1R mRNA expression in tumours. n = 8. (T-U) NPY protein levels in tumour tissues measured by human-specific ELISA kit, normalised to (T) total protein or (U) tissue weight, n = 8. (V-W) NPY protein levels in tumour tissues measured by mouse-specific ELISA kit, normalised to (V) total protein or (W) tissue weight, n = 8. (X) NPY mRNA expression in tumour tissues, n = 8. (Y) Plasma NPY levels at Day 0, Day 7, and Day 35, n = 4-5. Data are presented as mean ± SEM, and were analysed by one-way ANOVA with Tukey post hoc tests, \**p* < 0.05, \*\**p* < 0.01, \*\*\**p* < 0.001, \*\*\*\**p* < 0.0001.

In the Open Field Test (OFT) (Fig. 3B-F), the travel distance, number of entries and time spent in the central zone were reduced in mice in the Stress group (Fig. 3C-E, Ctrl vs Stress, *p* < 0.05), suggesting increased anxiety-like behaviour. BIBO treatment improved only the number of centre entries (Fig. 3D, Stress vs Stress+BIBO, *p* < 0.05). Total distance travelled was similar among groups (Fig. 3F). The performance in the Elevated Plus Maze (EPM) echoed the changes in the OFT (Fig. 3G,H), with reduced entries and time spent in open arms (Fig. 3G,H, *p* < 0.05) in the Stress group, which, however, was ameliorated by BIBO treatment (Fig. 3G,H). In the Tail Suspension Test (TST), mice from the Stress group exhibited prolonged immobile time, suggesting depression-like behaviour (*p* < 0.05 compared with the Ctrl), which was not affected by BIBO treatment (Fig. 3I). Tests outcomes collectively demonstrate that under tumour-bearing conditions, the chronic restraint induced sustained anxiety- and depression-like behaviours, while inhibiting NPY1R is only partially effective.

From tumour onset (day 15), chronic stress significantly accelerated tumour growth, which was effectively antagonised by BIBO (Fig. 3J, *p* < 0.001, Ctrl vs Stress, Stress vs Stress+BIBO). We next observed that stress significantly promoted tumour growth and tumour weight, which were markedly inhibited by BIBO treatment (Fig. 3K,L). Notably, body weight was not significantly different between groups (Fig. 3M). To further evaluate the effect of chronic stress on tumour behaviours, we used HGC27-luciferase cells and *in vivo* bioluminescence imaging. It appeared that stress significantly enhanced tumour growth, which was markedly inhibited by BIBO treatment (Fig. 3N, *p* < 0.01, Ctrl vs Stress; *p* < 0.0001, Stress vs stress+BIBO). Metastasis was not detected in distant organs in all group due to the short duration. Immunofluorescence staining showed that chronic stress significantly increased the proportion of KI67-positive cells and decreased cleaved caspase3-positive cells in tumour tissue, suggesting enhanced cell proliferation and reduced apoptosis, which both were abrogated by BIBO treatment (Fig. 3O, *p* < 0.01, Ctrl vs Stress, *p* < 0.001, Stress vs Stress+BIBO, Fig. 3P, *p* < 0.01, Ctrl vs Stress, *p* < 0.01, Stress vs Stress+BIBO).

Subsequently, we examined changes in NPY/NPY1R signalling in tumour tissues. Chronic stress significantly increased NPY1R level in tumour tissues, which was markedly inhibited by BIBO treatment (Fig. 3Q, *p* < 0.0001, Ctrl vs Stress; *p* < 0.0001, Stress vs Stress+BIBO). Changes in NPY1R were further validated at both protein and transcript levels (Fig. 3R,S). Chronic stress significantly promoted NPY release in the tumour, which was significantly inhibited by BIBO treatment when measured by a human ELISA kit (Fig. 3T, *p* < 0.0001, Ctrl vs Stress, *p* < 0.001, Stress vs Stress+BIBO; Fig. 3U, *p* < 0.001, Ctrl vs Stress, *p* < 0.01, Stress vs Stress+BIBO). However, when a mouse-specific ELISA kit was used, no differences in NPY levels were observed among groups (Fig. 3V,W). qPCR analysis further confirmed that chronic stress significantly upregulated NPY mRNA expression in tumour tissues, which again was normalised by BIBO treatment (Fig. 3X, *p* < 0.0001, Ctrl vs Stress, Stress vs Stress+BIBO). Plasma NPY levels were not changed at the different experimental time points (Day 0, Day 7, and Day 35; Fig. 3Y). Given that we used the human-derived HGC27 cell line, these results suggest that under chronic stress conditions, the increased NPY within tumours is primarily produced and released by the tumour cells, but not by host innervation.

To further validate the direct effects of NPY/NPY1R signalling on gastric cancer cells, we performed functional studies on HGC27 cells *in vitro* (Extended Data Fig.2). NPY or NPY1R agonist [Leu31, Pro34]-Neuropeptide Y (Leu) significantly increased the proportion of KI67-positive HGC27 cells, whereas NPY1R antagonist BIBO significantly reduced KI67-expressing cells number (Extended Data Fig.2A, *p* < 0.05, Ctrl vs NPY and Leu; *p* < 0.001, Ctrl vs BIBO). Wound healing assay showed that both NPY and Leu significantly promoted HGC27 cell migration at 12h, 24h, and 48h, while BIBO suppressed it (Extended Data Fig.2B-E, 12h: *p* < 0.001, Ctrl vs NPY, Ctrl vs Leu, *p* < 0.05, Ctrl vs BIBO; 24h: *p* < 0.001, Ctrl vs NPY, Ctrl vs Leu, *p* < 0.01, Ctrl vs BIBO; 48h: *p* < 0.001, Ctrl vs NPY, Ctrl vs Leu, Ctrl vs BIBO). NPY and Leu also enhanced the invasive behaviour of HGC27 cells, whereas BIBO significantly inhibited this process (Extended Data Fig.2F, *p* < 0.001, Ctrl vs NPY, Ctrl vs Leu, Ctrl vs BIBO). Furthermore, NPY and Leu treatments decreased the proportion of TUNEL-positive cells, whereas BIBO treatment reversed this change (Extended Data Fig.2G, *p* < 0.01, Ctrl vs NPY, Ctrl vs Leu, Ctrl vs BIBO).

To summarise, the results of *in vivo* and *in vitro* studies consistently demonstrate that chronic stress enhances the NPY/NPY1R levels within gastric tumour tissues and promotes cancer growth and tumour cell proliferation, whereas the NPY1R-specific inhibitor BIBO3304 can effectively suppress such stress-induced responses.

### 3. Under chronic stress, tumour-associated neuronal signals are significantly correlated with the central amygdala (CeA) activity

We next investigated the responsible brain nuclei, using a retrograde trans-multisynaptic tracing to systematically trace projection pathways from the tumour site to the central nervous system (CNS) in BALB/c-nude mice with subcutaneously inoculated HGC27 cells (Fig. 4A). After stable tumour formation (∼4 weeks), pseudorabies virus (PRV) carrying EGFP (PRV-EGFP, titer: 5 × 10C PFU/mL, 15 μL) was injected into three sites in the tumour. One week later, PRV-EGFP signals originating from the tumour site travelled through the spinal cord and reached the brain in both the control mice and mice with chronic stress groups. Specifically, PRV-EGFP signals were observed rostrally through the dorsal horn of the sacral, lumbar, thoracic, and cervical spinal cord (Extended Data Fig.3A,B), suggesting the polysynaptic connection pathways between the tumour and the CNS. Furthermore, under chronic stress, EGFP signals were most prominent in the CeA, followed by the dorsomedial hypothalamus (DMH) (Fig. 4B-E). In contrast, PRV-EGFP signals were not detected in the CeA and DMH in the Ctrl group, suggesting that chronic stress promoted the connection between the brain and tumour.

**Fig. 4.**
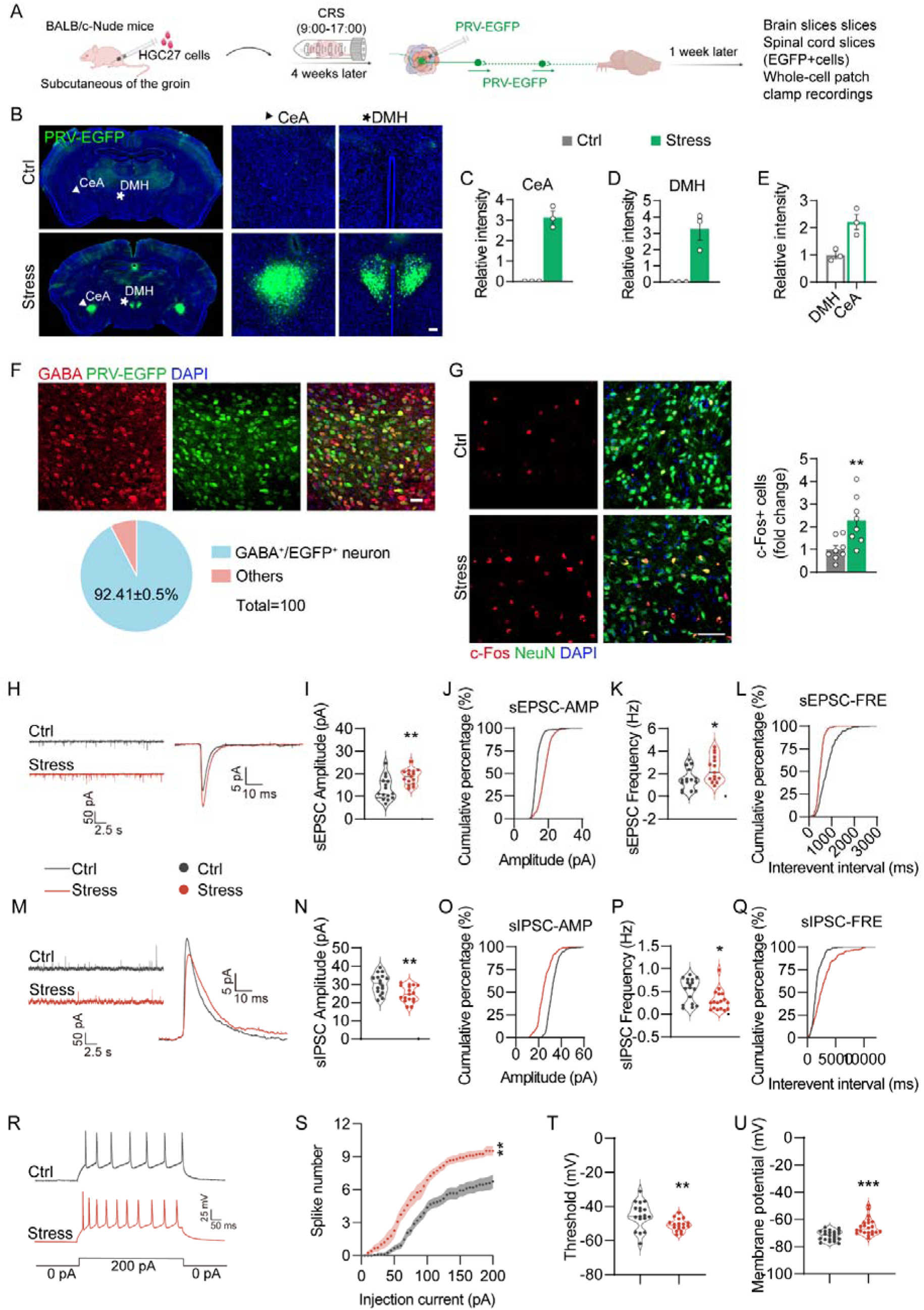
Chronic restraint stress triggers a brain-tumour axis by activating the central amygdala (CeA). (A) Schematic illustration of the experiments: retrograde multi-synaptic tracing of the tumour-brain axis using pseudorabies virus labelled by EGFP (PRV-EGFP). (B) Representative images of brain sections showing CeA, scale bar: 50 μm. PRV-EGFP intensity in the (C) CeA and (D) Dorsomedial hypothalamus (DMH) of ctrl tumour-bearing mice and mice with stress, n = 3. (E) Comparison of PRV-EGFP intensity in the CeA and DMH, n = 3. (F) Representative images of GABA immunostaining in the CeA, scale bar: 50 μm. Below: Quantification of PRV-EGFP and GABA double-positive cells, n = 4. (G) Representative images of c-Fos and NeuN immunostaining in the CeA of ctrl tumour-bearing mice and mice with stress, scale bar: 50 µm. Right: Quantification of c-Fos-positive cells relative to the number of NeuN-positive cells. n = 8. (H) Representative sEPSC traces and curves in the CeA of ctrl tumour-bearing mice and mice with stress. (I) Quantification of spontaneous excitatory postsynaptic currents (sEPSC) amplitude, n = 15 cells, n = 3 mice. (J) Cumulative distribution of sEPSC amplitude. (K) sEPSC frequency, n = 15 cells, n = 3 mice. (L) Cumulative distribution of sEPSC frequency. (M) Representative images of sIPSC traces and curve in the CeA of ctrl tumour-bearing mice and mice with stress. (N) Quantification of spontaneous inhibitory postsynaptic current (sIPSC) amplitude, n = 17 cells, n = 3 mice. (O) Cumulative distribution of sIPSC amplitude. (P) Quantification of sIPSC frequency in the CeA. n = 17 cells, n = 3 mice. (Q) Cumulative distribution of sIPSC frequency. (R) Representative action potential firing traces in the CeA of ctrl tumour-bearing and mice with stress. (S) Total spike counts of CeA neurons, n = 15 cells, n = 3 mice. (T) Action potential threshold of CeA neurons, n = 18 cells, n = 3 mice. (U) Resting membrane potential of CeA neurons, n = 22 cells, n = 3. Data are presented as mean ± SEM, and were analysed by two-tailed *t* tests, \**p* < 0.05, \*\**p* < 0.01, \*\*\**p* < 0.001.

Given the central role of the CeA in processing stress and negative affect, as well as the prominent PRV-EGFP signal in the CeA, we first characterised the types of neurons labelled with PRV-EGFP. Over 90% (Fig. 4F, 92.41 ± 0.5%) of PRV-EGFP-positive neurons were GABAergic interneurons. To verify the activation status of the CeA neurons, c-Fos staining was used. The number of c-Fos-positive cells in the CeA was significantly increased in the tumour-bearing mice with stress (Fig. 4G, *p* < 0.01). Next, we used whole-cell patch clamp to assess the function of CeA neurons (Fig. 4H). The amplitude of spontaneous excitatory postsynaptic currents (sEPSC) of CeA neurons in chronically stressed tumour-bearing mice was significantly enhanced in the tumour-bearing mice with stress when compared to the Ctrl group (Fig. 4I, *p* < 0.01), and the cumulative distribution curve of amplitude was significantly shifted to the right (Fig. 4J). The sEPSC frequency was also significantly increased (Fig. 4K, *p* < 0.05), and the cumulative distribution curve of frequency was shifted to the left (Fig. 4L), suggesting increased excitatory synaptic input. At the same time, the amplitude of spontaneous inhibitory postsynaptic currents (sIPSC) in CeA neurons (Fig. 4M) was significantly reduced in the tumour-bearing mice with stress (Fig. 4N, *p* < 0.01), whereas the cumulative distribution curve of amplitude was shifted to the left (Fig. 4O). Moreover, the sIPSC frequency was also significantly decreased (Fig. 4P, *p* < 0.05), and the cumulative distribution curve of frequency was shifted to the right (Fig. 4Q), suggesting reduced inhibitory synaptic input. The total count of CeA neuronal spikes was significantly increased in the tumour-bearing mice with stress (Fig. 4R,S, *p* < 0.01), while the action potential threshold was significantly decreased (Fig. 4T, *p* < 0.01), while the resting membrane potential was significantly depolarised (Fig. 4U, *p* < 0.001), indicating that the overall excitability of CeA neurons is enhanced under chronic stress conditions. Together, under chronic stress conditions, the CeA neuronal excitability and activity were significantly higher in tumour-bearing mice, suggesting that CeA activation may serve as a central neural mechanism linking emotional disturbances with tumour progression.

### 4. Chronic stress enhanced CeA-driven sympathetic overflow, increasing NE release and stimulating NPY synthesis in tumour cells, thereby facilitating gastric cancer progression

CeA regulate sympathetic output through the medulla oblongata and the intermediolateral nucleus (IML) of the spinal cord ^41,42^. We hypothesised that CeA neuron activation mediates stress-induced gastric tumour progression by enhancing peripheral sympathetic activity. To test this hypothesis, we examined the levels of NE, Ach, and cortisol in both tumour tissues and plasma (Fig. 5A-C, Extended Data Fig.4A-F). NE levels in the tumour tissues and serum were significantly elevated in the Stress group (Fig. 5A,B, *p* < 0.0001 vs Ctrl); however, there was no difference between the Stress+BIBO and the Stress groups. The NE ratio between tumour tissue and plasma was also significantly increased in the Stress group (Fig. 5C, *p* < 0.01), with no change between the Stress+BIBO and the Stress groups. In contrast, Ach levels were similar between groups (Extended Data Fig.4A-C). Cortisol levels in both tumour tissue and plasma were significantly elevated in the Stress group (Extended Data Fig.4D,E, both *p* < 0.0001), but no significant difference was found between the Stress+BIBO group and the Stress group; neither the tumour tissue/plasma cortisol ratio changed (Extended Data Fig.4F). These results suggest chronic stress mainly promotes sympathetic neurotransmitter NE release in the tumour tissues, without affecting vagal neurotransmitter Ach and the stress hormone cortisol.

**Fig. 5.**
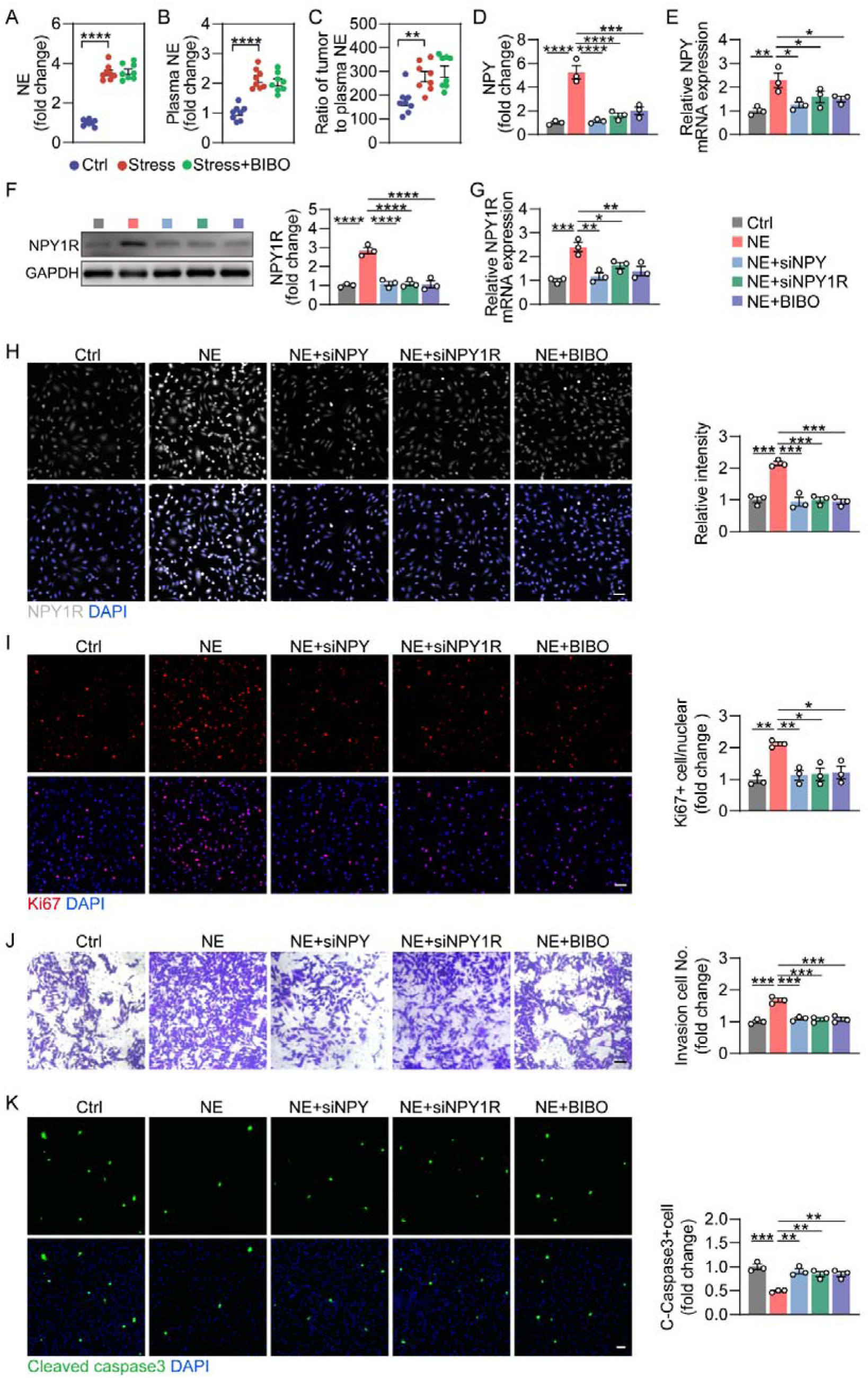
Chronic stress increased the release of the sympathetic neurotransmitter, which was inhibited by BIBO inhibitors. (A) Norepinephrine (NE) levels in tumour tissues of the Ctrl, Stress, and Stress+BIBO groups, n = 8. (B) NE levels in the plasma of the Ctrl, Stress, and Stress+BIBO groups, n = 8. (C) Tumour tissue/plasma NE ratio in the Ctrl, Stress, and Stress+BIBO groups, n = 8. (D) NPY secretion, (E) NPY mRNA expression, (G) NPY1R mRNA expression, (F) NPY1R protein levels, (H) representative images of NPY1R staining (scale bar: 50 μm) in HGC27 cells treated with NE, NE+siNPY, NE+siNPY1R, or NE+BIBO, n = 3 independent experiments. Right: quantification of NPY1R intensity. n = 3 independent experiments. (I-K) Functional assays in HGC27 cells under the same conditions in (D): (I) proliferation, (J) invasion, and (K) apoptosis, scale bar: 50 μm, n = 3 independent experiments. Data are presented as mean ± SEM, and were analysed by one-way ANOVA with Tukey’s post hoc tests, \**p* < 0.05, \*\**p* < 0.01, \*\*\**p* < 0.001, \*\*\*\**p* < 0.0001.

To further verify whether intratumoral NE can directly induce NPY/NPY1R production in gastric cancer cells, HGC27 cells were treated with 10 μmol/L NE ^43^. NPY/NPY1R signalling was manipulated by siNPY, siNPY1R, and BIBO, thus necessitating 5 groups: Ctrl, NE, NE+siNPY, NE+siNPY1R, NE+BIBO. NPY secretion, NPY/NPY1R expression, and associated cellular functional changes were systematically assessed (Fig. 5D-K). Small interfering RNAs (siRNAs) targeting NPY, NPY1R significantly downregulated the mRNA expression of the corresponding genes in gastric cancer cells (Extended Data Fig.4G,H). Compared with the Ctrl group, NPY levels in the culture medium were significantly increased after NE treatment (Fig. 5D, *p* < 0.0001), whereas NE-induced increases in NPY secretion (Fig. 5D) and NPY mRNA expression (Fig. 5E) were significantly inhibited by siNPY, siNPY1R, and the NPY1R-specific inhibitor BIBO (Fig. 5D,E). NE also significantly increased NPY1R protein levels and mRNA expression in HGC27 cells, which were effectively inhibited by siNPY, siNPY1R, and BIBO (Fig. 5F-H). In addition, NE treatment significantly increased cell proliferation (Fig. 5I, *p* < 0.01, Ctrl vs NE) and invasion (Fig. 5J, *p* < 0.001, Ctrl vs NE) and reduced the apoptosis (Fig. 5K, *p* < 0.001, Ctrl vs NE) in HGC27 cells, which were effectively suppressed by siNPY, siNPY1R, and BIBO (Fig. 5I). These results suggest that NE can directly promote the synthesis and release of NPY and NPY1R production in cancer cells, associated with enhanced gastric cancer cell proliferation and invasion.

We next investigated the downstream pathways of NPY/NPY1R, focussing on MAPK/Erk1/2 signalling (Extended Data Fig.5), which has been involved in the progression of other cancers, e.g., prostate cancer and breast cancer ^44,45^. In the Stress group tumour tissues, p-Erk1/2 level was significantly increased, which was inhibited by BIBO (Extended Data Fig.5A, *p* < 0.0001, Ctrl vs Stress, Stress vs Stress+BIBO). In HGC27 cells, NPY or the NPY1R agonist Leu significantly upregulated p-Erk1/2 levels, which were significantly reduced by BIBO (Extended Data Fig.5B, *p* < 0.001, Ctrl vs NPY, Ctrl vs Leu, Ctrl vs BIBO). Furthermore, in the HGC27 cells, NE treatment also increased p-ERK1/2 levels, which were reversed by siNPY, siNPY1R, as well as the BIBO (Extended Data Fig.5C). In gastric cancer patient samples, p-Erk1/2 levels were significantly higher than those in the paracancerous tissues (Extended Data Fig.5D, *p* < 0.01).

Overall, chronic stress increases sympathetic neurotransmitter NE release in gastric tumours, which activates NPY/NPY1R signalling through MAPK/Erk1/2 pathway, thereby facilitating tumour progression.

### 5. Central intervention - targeting CeA activity to modulate tumour progression

Given tumour-specific response of CeA to stress, we further investigated whether manipulating with CeA neuronal activity can alter stress-related tumour progression (Fig. 6A) by using the Designer Receptors Exclusively Activated by Designer Drug (DREADD) system: excitatory AAV-hM3D(Gq), or inhibitory AAV-hM4D(Gi) injected bilaterally into the CeA region. This yielded 4 groups: Ctrl (tumour), Stress (tumour+stress), hM3D(Gq)(tumour+CeA activation), and Stress+hM4D(Gi) (tumour+stress+CeA inhibition). AAV infection was mainly confined to the CeA region (Fig. 6B). After CNO administration, neuronal firing activity in the hM3D(Gq) group was significantly enhanced, whereas that in the hM4D(Gi) group was markedly suppressed (Fig. 6C), suggesting effective regulation of CeA activity. Compared with the Ctrl tumour-bearing group, c-Fos level in the CeA neurons was significantly increased in the Stress and hM3D(Gq) groups, which was normalised in the hM4D(Gi) group (Fig. 6D, *p* < 0.01, Ctrl vs Stress and hM3D(Gq); *p* < 0.05, Stress vs Stress+hM4D(Gi)). To evaluate the effects on cognitive behaviours, OFT and TST were performed. In the OFT, travel distance, central entries, and time spent in the centre zone were significantly reduced in the Stress group (Fig. 6E-G, all *p* < 0.05), which was not changed in the hM3D(Gq) and Stress+hM4D(Gi) groups compared with the Ctrl group (Fig. 6E-H, all *p* < 0.05). During TST, immobility time was significantly prolonged in the Stress group, hM3D(Gq), and Stress+hM4D(Gi) groups (Fig. 6J, *p* < 0.05, Ctrl vs Stress and hM3D(Gq)). These results suggest that under tumour-bearing conditions, CeA neuronal activity primarily regulates anxiety-like behaviour.

**Fig. 6.**
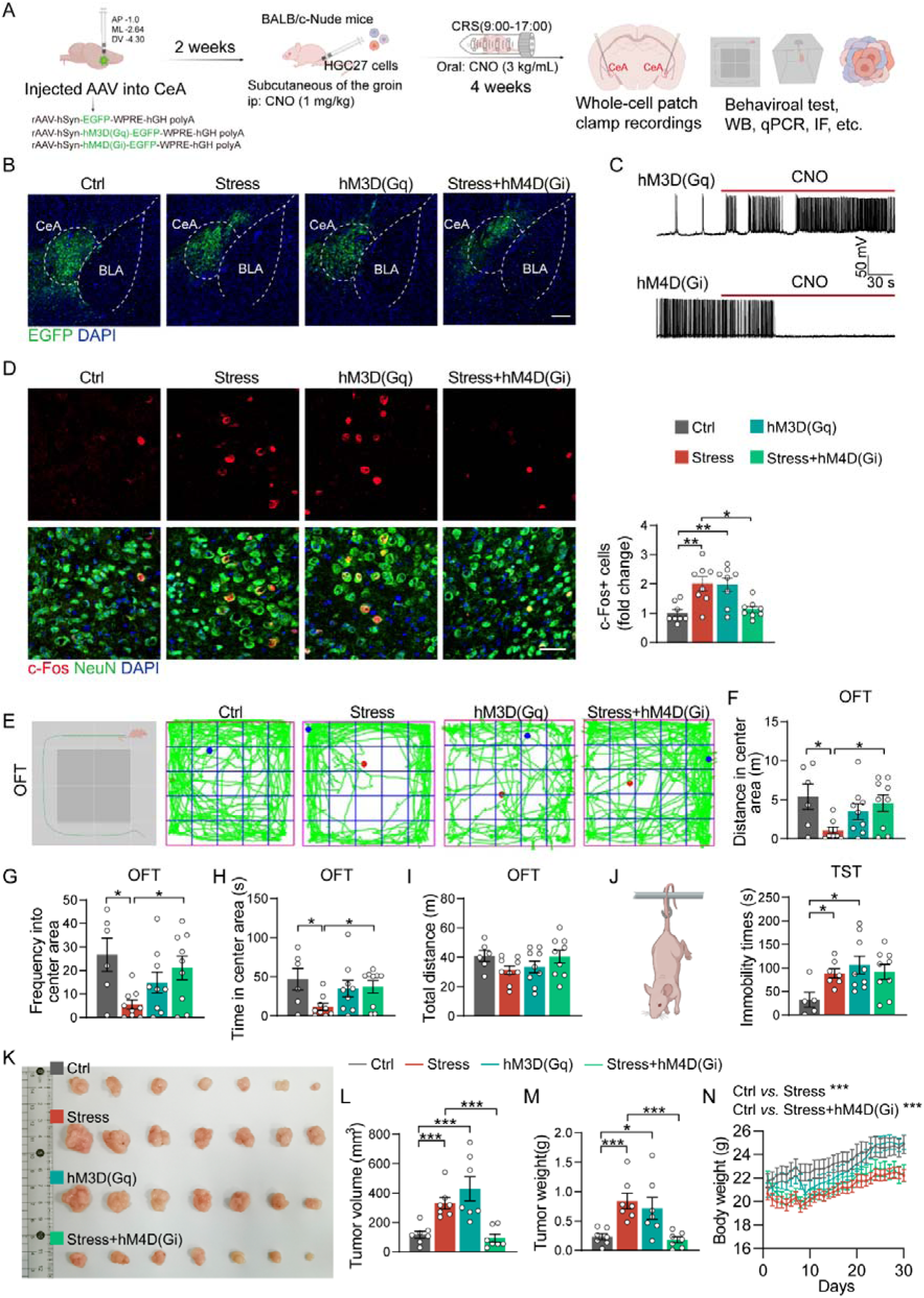
Chemogenetic manipulation of central amygdala (CeA) activity to regulate gastric tumour growth. (A) Schematic diagram of experimental design. (B) Representative images of AAV infection in the CeA. Scale bar: 200 μm. (C) Whole-cell patch-clamp recordings showing neuronal firing activity in CeA neurons after CNO administration. hM3D(Gq) group exhibited significantly increased firing, while hM4D(Gi) group showed markedly suppressed firing. (D) Representative images of c-Fos and NeuN immunostaining in CeA neurons. Scale bar: 50 μm. Right: quantification of c-Fos intensity, n = 8. (E-H) Open field test (OFT) to assess anxiety-like behaviour. (E) Representative movement traces; (F) distance travelled, (G) frequency of entries, and (H) time spent in the centre, n = 6-9 mice. (I) Total distance travelled in OFT, n = 6-9 mice. (J) Immobility time during the tail suspension test (TST), n = 5-9 mice. (K) Representative images of tumours at the endpoint. (L) Tumour volume and (M) tumour weight at endpoint, n = 7 mice. (N) Changes in body weight over time, n = 10-11 mice. Data are presented as mean ± SEM, and were analysed by one-way ANOVA with Tukey’s post hoc test, \**p* < 0.05, \*\**p* < 0.01, \*\*\**p* < 0.001.

Subsequently, we evaluated the effects of CeA neuronal activity on tumour growth. Both tumour volume and weight were significantly increased in the Stress group when compared to the Ctrl group (Fig. 6L,M, both *p* < 0.001). Activation of CeA neurones alone due to hM3D(Gq) significantly promoted tumour growth to a similar tumour volume (Fig. 6L, *p* < 0.001) and weight (Fig. 6M, *p* < 0.05) as in the Stress group; conversely inhibition of neuronal activity in CeA by hM4D(Gi) significantly suppressed stress-induced tumour growth (Fig. 6L,M, *p* < 0.001 vs Stress and hM3D(Gq) groups). In addition, weight loss was induced by long-term stress, but not by modifying CeA neuronal activity (Fig. 6N, *p* < 0.001, Ctrl vs Stress, Ctrl vs Stress+hM4D(Gi)).

To further dissect how CeA manipulation affects the tumour microenvironment, we examined markers of cell proliferation and apoptosis, as well as the NPY/NPY1R axis, and the MAPK signalling pathway in tumour tissue, (Fig. 7). First, Ki67 levels in tumour tissues were significantly increased in the Stress and hM3D (Gq) groups; the Ki67 was rescued to the Ctrl level in the Stress+hM4D(Gi) group (Fig. 7A, *p* < 0.01, Ctrl vs Stress, Ctrl vs hM3D(Gq); *p* < 0.05, Stress vs Stress+hM4D(Gi)). Correspondingly, Cleaved caspase-3 levels in tumour tissues were significantly reduced in the Stress and hM3D(Gq) groups, which was partially restored in the Stress+hM4D(Gi) group (Fig. 7B, *p* < 0.0001, Ctrl vs Stress, Ctrl vs hM3D(Gq), Stress vs Stress+hM4D(Gi)). Given that increased NPY in cancer may originate from enhanced sympathetic activity, we further examined NE in the tumour tissue. CeA activation (hM3D(Gq)) significantly increased NE levels in tumour; the NE was significantly reduced by inhibiting CeA neuronal activity (Fig. 7C, *p* < 0.0001, Ctrl vs Stress, Ctrl vs hM3D(Gq), Stress vs Stress + hM4D(Gi)). The changes in plasma NE levels (Fig. 7D, *p* < 0.0001, Ctrl vs Stress, Ctrl vs hM3D(Gq), Stress vs Stress+hM4D(Gi)) and ratio of NE in the tumour tissue/plasma (Fig. 7E, *p* < 0.001, Ctrl vs Stress, *p* < 0.05, Ctrl vs hM3D(Gq), *p* < 0.01, Stress vs Stress+hM4D(Gi)) mirrored those in tumour tissues. In contrast, there were no significant differences in Ach levels or their ratios of tumour tissues/plasma among groups (Extended Data Fig.6A-C). Stress increased cortisol levels in both tumour tissues and plasma (Extended Data Fig.6D,E, *p* < 0.0001). Notably, inhibition of CeA neurones doubled cortisol levels in both tumour tissue and serum (Extended Data Fig.6D,E). The ratio of cortisol in tumour tissue/plasma was similar among groups (Extended Data Fig.6F). Furthermore, NPY1R protein levels measured by immunostaining and Western blotting, NPY1R mRNA expression, NPY concentration and NPY mRNA expression within tumour tissues were consistently increased in the Stress and hM3D(Gq) groups, but were reduced to normal values by inhibition of CeA neuronal activity in the Stress+hM4D(Gi) group (Fig. 7F-J). There were no differences in plasma NPY levels among groups (Fig. 7K). In addition, NPY downstream signalling MAPK related p-Erk1/2 levels were also increased by both stress and activation of CeA neurones alone; the p-Erk1/2 was reduced to normal values by CeA neuronal inhibition in tumour-bearing mice with stress (Fig. 7L, *p* < 0.05, Ctrl vs Stress, Ctrl vs hM3D(Gq), Stress vs Stress+hM4D(Gi)).

**Fig. 7.**
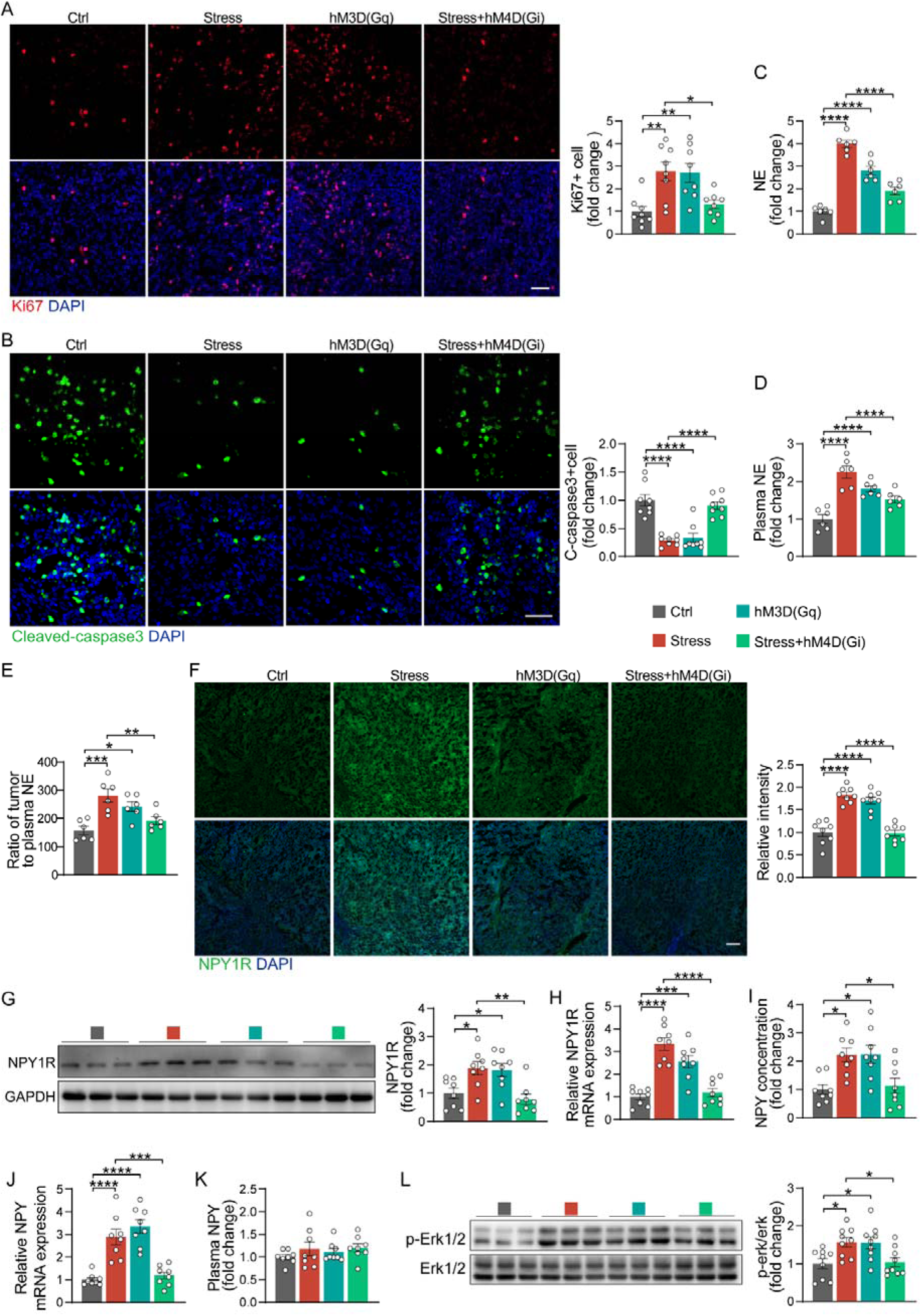
CeA neuronal activities regulate tumour proliferation and apoptosis via NE-NPY/NPY1R-MAPK signalling in tumours. (A) Representative images of Ki67 immunostaining in tumours of Ctrl, Stress, hM3D(Gq), and Stress+hM4D (Gi) tumour-bearing groups, scale bar: 50 μm. Right: Quantification of Ki67-positive cells, n = 8. (B) Representative images of Cleaved caspase-3 immunostaining in tumours, scale bar: 50 μm. Right: Quantification of Cleaved caspase-3-positive cells, n = 8. (C) NE levels in tumour tissues and (D) the plasma, n = 6. (D) Ratio of NE levels in the tumour tissue/plasma, n = 6. (F) Representative images of NPY1R immunofluorescence staining in the tumours, scale bar: 50 μm. Right: Quantification of NPY1R fluorescence intensity, n = 8. Statistical test: one-way ANOVA with Tukey’s post hoc test. (G) NPY1R protein and (H) mRNA levels in tumours, n = 8. (I) NPY protein and (J) mRNA levels in tumour tissues, n = 8. (K) NPY levels in the plasma, n = 8. (L) p-Erk1/2 protein levels in tumour tissues, n = 8. Data are presented as mean ± SEM, and were analysed by one-way ANOVA with Tukey’s post hoc test, \**p* < 0.05, \*\**p* < 0.01, \*\*\**p* < 0.001, \*\*\*\**p* < 0.0001.

These results indicate that CeA activity plays a key role in stress-related acceleration of gastric tumour growth, driven by intratumoral sympathetic activity and associated NPY/NPY1R–MAPK/ Erk1/2 signalling pathway.

## Discussion

Cancer progression is a dynamic process shaped by interactions within the tumour microenvironment and between the tumour and the nervous system ^46–48^, yet how brain circuits engaged by emotional stress influence tumour behaviour remains largely unknown. Here, we define a centrally driven brain-tumour axis in gastric cancer, in which chronic stress engages the CeA as a key neural hub that translates emotional stress into heightened peripheral sympathetic output. This process actively remodels the tumour microenvironment through activation of the NPY/NPY1R/MAPK–Erk1/2 signalling cascade, thereby accelerating tumour growth. Our findings establish a direct causal link between a discrete emotionalCprocessing neural circuit and tumour progression, extending the concept of the brain-tumour axis beyond associative effects of stress to a mechanistically defined and potentially modifiable neural pathway (Fig 8).

**Fig. 8.**
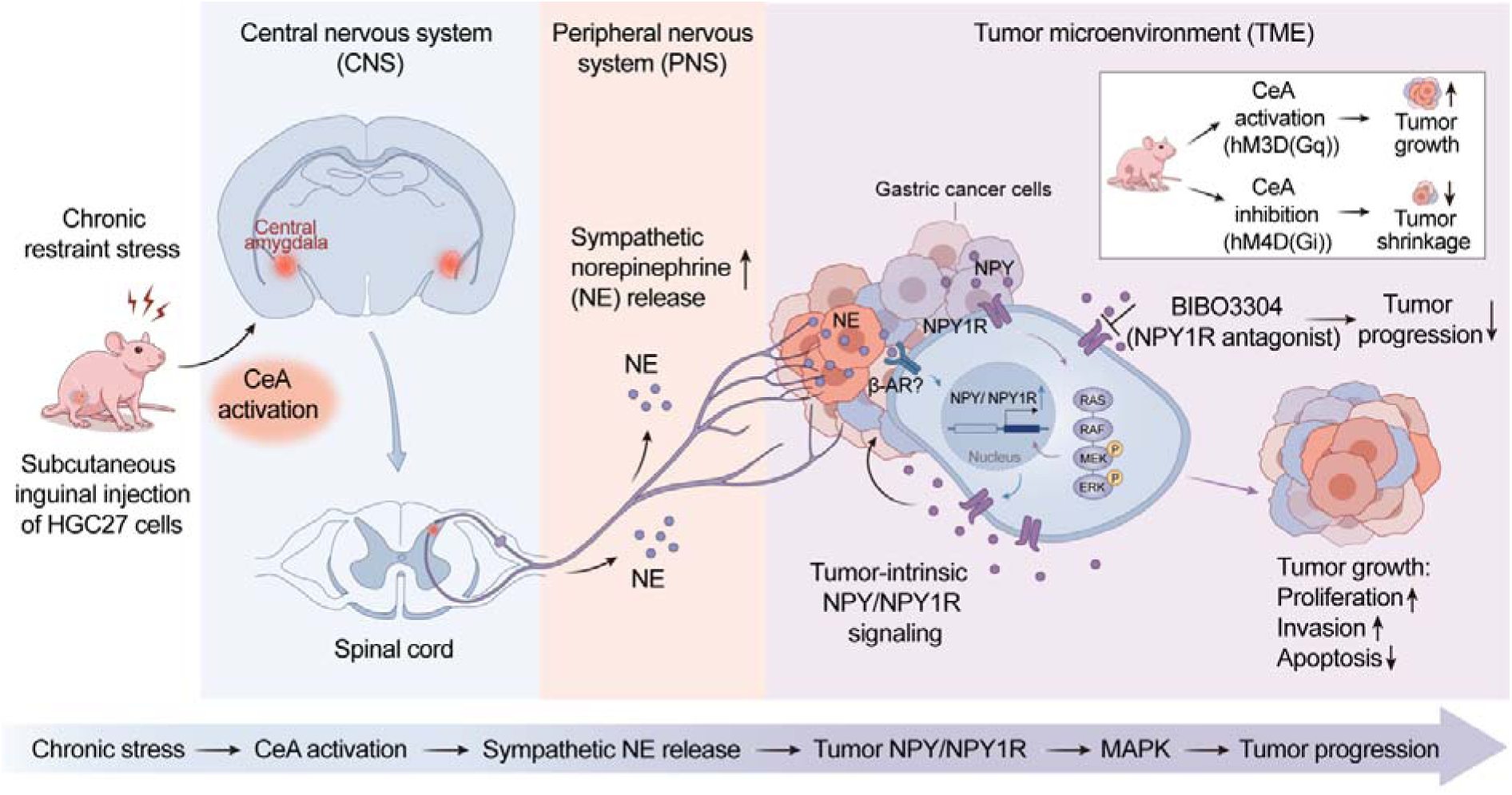
Chronic stress drives gastric cancer progression through a CeA-tumour axis engaging NPY signalling. Chronic stress activates CeA, leading to sympathetic nerve infiltration and increased release of NE within the tumour microenvironment. NE, in turn, upregulates both NPY and its receptor NPY1R in cancer tissue by an autocrine/paracrine signalling loop. The binding of NPY to NPY1R subsequently triggers the MAPK/ERK1/2 pathway, thereby promoting tumour cell proliferation and invasion. Consequently, chronic stress-associated efferent neural signals drive gastric cancer progression. Our study defines a malignant brain-tumour axis, wherein CeA-originated stress signalling enhances tumour growth through the NE-NPY/NPY1R-MAPK/Erk1/2 cascade.

Recent advances in cancer neuroscience have highlighted the importance of the bidirectional brain-gut axis communication in gastrointestinal tumour biology, shifting the field away from purely systemic neuroendocrine models towards circuit-based regulation of tumour progression ^49^. However, although psychological stress and mental health states are increasingly recognised to shape gastric cancer outcomes, largely through elevated circulating catecholamines and glucocorticoids^26,50^, how central emotional circuits directly and causally influence peripheral tumours has remained unclear.

Classical autonomic neurotransmitters, such as NE ^51^, Ach^52^, have been implicated in gastric cancer progression, have been associated with gastric cancer progression; however, these observations do not resolve how central stress processing is converted into tumour-specific autonomic signalling. In this study, we demonstrate that the CeA, a key node in adverse emotional processing, functionally links psychological stress to gastric tumour progression. Chemogenetic gain- and loss-of-function approaches reveal that CeA neuronal activity is both necessary and sufficient to modulate stress-associated tumour growth, identifying the CeA as an upstream regulator of pro-tumourigenic autonomic output.

A major discovery of this study is that the brain regulation of tumour progression is mediated by the sympathetic nervous activity in the gastric tumour tissue, involving NPY/NPYR1 signalling. Chronic stress primarily increases catecholamine release from the sympathetic projections ^25,53^, demonstrated by increased sympathetic nerve infiltration and intratumoral NE level in stressed mice in this study. While NE is known to promote tumour progression through βCadrenergic signalling ^54,55^, our findings define a previously unrecognised mechanistic pathway in which NPY/NPY1R signalling links CeACmediated stress processing to peripheral sympathetic activation and gastric cancer progression. It supports dynamic neuro-tumour interactions in the tumour microenvironment ^56^. Consistent with our experimental findings, TCGA analysis performed in this study revealed increased NPY1R expression in stomach adenocarcinoma, which correlated positively with disease stage and inversely with patient survival. Our woundChealing assays in gastric cancer cells align with and extend prior findings in pancreatic cancer, where NPY/NPY1R signalling drives metastatic behaviour in a p53□dependent manner, supporting a conserved pro□migratory and pro□invasive role for this pathway across gastrointestinal malignancies ^57^. Although NPY is classically recognised as a sympathetic co□transmitter, our data indicate that under chronic stress conditions, NPY is predominantly synthesised and secreted by tumour cells in response to sympathetic NE signalling, functioning in an autocrine/paracrine manner to amplify NPY1R activation. Increased local NPY signalling is regulated by elevated neuronal activity in CeA under chronic stress. Local NPY availability also determines NPY1R expression in the tumour cells by a positive feedback loop. Activation of MAPK/ERK pathway has been consistently observed in several cancer studies ^58,59^. This study further confirms that NPY1R-dependent sympathetic activation, triggered by CeA response to chronic stress with tumour presence, acts as a unique upstream regulator required for tumour proliferation and potentially metastasis over time. Although our data identify MAPK/Erk1/2 as a critical downstream effector of NPY1R activation in gastric cancer, additional GPCR□linked pathways may operate in parallel and contribute to tumour progression under chronic stress.

Although vagal signalling has previously been implicated in gastric tumourigenesis, particularly via muscarinic receptor pathways, we did not observe changes in intratumoral acetylcholine levels under chronic stress. While this suggests that stress□induced tumour acceleration in our model is dominated by sympathetic rather than parasympathetic outputs, it does not preclude parallel modulation of muscarinic receptor signalling or downstream pathways independent of bulk acetylcholine concentration.

From a translational perspective, this brain-tumour axis offers multiple therapeutic entry points. Targeting the CeA may provide a novel strategy to suppress tumour progression, potentially through counselling, pharmacological modulation, or non-invasive neuromodulation such as repetitive transcranial magnetic stimulation ^60^. At the peripheral level, our findings suggest that NPY1R serves as a critical interface coupling neuronal-derived signals to canonical oncogenic pathways, which can also serve as a plausible cancer therapeutic target. Therefore, direct targeting of NPY1R using antagonists, such as BIBO3304, may represent a strategy for cancer patients suffering from anxiety disorders. Its limited blood-brain barrier permeability reduces potential central side effects ^61^. These approaches may be considered for adjunctive therapy with existing chemo- or immunotherapies.

Several limitations should be acknowledged. First, this study primarily examines tumour progression kinetics in chronically stressed, tumour□bearing mice; future work should assess whether combining standard chemo□or immunotherapies with NPY1R antagonism can further reduce tumour burden. Second, our mechanistic analyses were conducted using a single gastric cancer cell line, and validation across additional molecular subtypes will be required to establish broader relevance. Third, as NPY1R is a G□protein□coupled receptor engaging multiple downstream signalling cascades, future studies should delineate the full signalling network beyond MAPK/Erk1/2.

In summary, this study defines a multilevel regulatory axis linking chronic stress to accelerated gastric cancer progression, spanning CeA activity, peripheral sympathetic nervous outputs, neurotransmitter NPY release, as well as the increase in NPY1R and activation of its downstream signalling MAPK element Erk1/2 (Fig 8). The integration of systems neuroscience with cancer biology provides a mechanistic framework for understanding how psychological stress promotes gastric tumour progression, suggesting the brain-tumour axis as a new therapeutic target for improving the management of gastric cancer.

## Methods

### Human specimens

Human gastric adenocarcinoma tissues and adjacent non-tumour tissues were obtained from patients undergoing surgical resection at the Seventh Affiliated Hospital of Sun Yat-sen University (Shenzhen, China). All procedures were approved by the institutional ethics committee (Approval#: KY-2025-028-01), and written informed consent was obtained from all participants. Tumour staging was defined according to the AJCC TNM classification system. Fresh tissues for RNA and protein analyses were snap-frozen in liquid nitrogen and stored at -80□°C. Samples for histological analyses were fixed in 4% paraformaldehyde (PFA). Blood samples were collected from cancer patients (pre- and post-surgery) and healthy donors for plasma-based assays. Detailed information on human specimens is in Supplementary Table 1.

### Animals

All animal experiments were approved by the Institutional Animal Care and Use Committee of Sun Yat-sen University (Approval# SYSU-IACUC-2025-000353). All animal experiments were designed, conducted, and reported in accordance with the ARRIVE (Animal Research: Reporting of In Vivo Experiments) guidelines. Five-week-old male and female BALB/c nude mice were housed in a pathogen-free facility under 22 ± 1 °C, 50 ± 10% humidity, and 12 h light/12 h dark cycles, with ad libitum access to standard rodent chow and water. All mice were purchased from Chongqing Enswell Laboratory Animal Sales Co., Ltd and were randomly assigned to experimental groups. Investigators were blinded to group allocation during data acquisition and analysis when feasible.

### Cells

The human gastric cancer cell line HGC27 was purchased from Wuhan Procell Life Science & Technology Co., Ltd. (CL-0107). HGC27-luciferase cells were constructed by OBiO Technology (Shanghai) Corp., Ltd. HGC27 cells were maintained in RPMI-1640 (Gibco, 11875093) medium supplemented with 10% fetal bovine serum (Gibco, 10099141(C)) and 1% penicillin-streptomycin (Yuanpei Biotechnology (Shanghai), S110JV) at 37□°C in a humidified incubator with 5% CO□.

### Chronic restraint stress and xenograft procedure

Mice were randomly divided into three groups: Ctrl (tumour+vehicle), Stress (restraint+tumour+vehicle), and Stress+BIBO3304. Chronic restraint was initiated 7 days prior to tumour implantation and performed daily for 35 consecutive days (8□h per day). Subcutaneous xenografts were established by injecting HGC27 cells into the inguinal region of nude mice. Tumour size was measured once every 2 days using callipers, and volume was calculated as V = ½ × a × b². BIBO3304 (5□mg/kg) was administered intraperitoneally once every 2 days starting on the day of tumour implantation. Blood samples were collected at Days 0, 7 and 35 of the study, from the orbital veins.At the endpoint, animals were perfused with PBS after deep anesthesia with a sterile solution of 1.25% tribromoethanol (Avertin) in tertiary amyl alcohol/saline (250 mg/kg, i.p.; Nanjing Aibei Bio., China, M2960), and tissues were harvested for molecular and histological analyses.

### Chemogenetic manipulation of CeA neurons

AAV-delivered chemogenetic approaches were used to manipulate neuronal activity in the CeA. Mice received bilateral stereotaxic injections of AAV constructs expressing EGFP (Ctrl, rAAV-hSyn-EGFP-WPRE-hGH polyA), hM3Dq (excitatory DREADD, rAAV-hSyn-hM3D(Gq)-EGFP-WPRE-hGH polyA), or hM4Di (inhibitory DREADD, rAAV-hSyn-hM4D(Gi)-EGFP-WPRE-hGH polyA). Stereotaxic coordinates were: AP -1.0 mm, ML ± 2.64 mm, DV - 4.30 mm (relative to bregma). Following viral expression, clozapine-N-oxide (CNO, 1□mg/kg, intraperitoneal) was administered to activate DREADDs, followed by maintenance delivery in drinking water.

### PRV-mediated retrograde trans-synaptic tracing

To map tumour-brain connectivity, PRV-CAG-EGFP was injected into established tumours. Virus (5 × 10□PFU/mL) was diluted and injected into the tumour tissue. Animals were perfused 4 -10 days post-injection. Brain and spinal cord tissues were collected, fixed, cryoprotected, and sectioned at 50 μm thickness. EGFP-labelled neurons were visualised using whole-slide fluorescence imaging, and distribution patterns were analysed.

### Brain slice recording

Mice were decapitated after being anesthetised with a sterile solution of 1.25% tribromoethanol (Avertin) in tertiary amyl alcohol/saline (250 mg/kg, i.p.; Nanjing Aibei Bio., China, M2960), and then the brain was placed in a Vibroslice (VT 1200S; Leica) to prepare coronal amygdala slices (300 μm) in ice-cold artificial cerebral spinal fluid (ACSF: 195 mM sucrose, 2.0 mM KCl, 0.2 mM CaCl□, 12 mM MgSO□, 1.3 mM NaH□PO□, 26 mM NaHCO□, and 10 mM glucose). Slices were incubated for 30 min at 34°C with recording ACSF saturated with a 95% O□and 5% CO□mixture. After incubation, brain slices were transferred to the recording chamber, which was continuously perfused at 3 mL/min with recording ACSF maintained at 32 - 34 °C. The recording ACSF contained: 126 mM NaCl, 3.0 mM KCl, 1.25 mM NaH□PO□·H□O, 2.0 mM CaCl□·H□O, 1.0 mM MgSO□, 26 mM NaHCO□, and 10 mM glucose. All solutions were continuously bubbled with a 95% O□and 5% CO□mixture. For whole-cell recording, pipettes were filled with an internal solution (126 mM K-gluconate, 2.0 mM KCl, 10 mM HEPES, 2.0 mM MgCl□, 4.0 mM Na□ATP, 0.4 mM Na□GTP, and 10 mM phosphocreatine (pH 7.25)). Spontaneous excitatory postsynaptic currents (sEPSCs) were recorded under voltage clamp at - 70 mV, while spontaneous inhibitory postsynaptic currents (sIPSCs) were recorded at 0 mV. For chemogenetic manipulations, after stabilising the recording of membrane potential and spike activity, the Designer Receptors Exclusively Activated by Designer Drugs (DREADD) agonist Clozapine N-oxide (CNO, 5 μM) was bath-applied, followed by continuous recording of membrane potential and spike activity for 3 min. All recordings were performed using conventional patch-clamp amplifiers [EPC10 (HEKA, Germany)] and analysed with ClampFit 11.2 (Molecular Devices, CA, USA). Data were collected only when the initial series resistance was within 20 - 30 MΩ, with signals sampled at 10 kHz and filtered at 1 kHz.

### Behavioural assays

All behavioural experiments were conducted in a dedicated behavioural testing room under controlled environmental conditions (22 ± 2 °C; 50-60% humidity) with minimal noise and olfactory interference. Mice were acclimated to the testing environment for at least 30 min prior to each experiment. All assays were performed during the same circadian phase to minimise variability. Apparatuses were cleaned with 75% ethanol between trials. Behavioural data were acquired using an automated video-tracking system and verified by blinded manual scoring.

### Elevated plus maze (EPM)

The elevated plus maze consisted of two open arms and two enclosed arms arranged in a cross configuration and elevated above the floor. Each mouse was placed in the central platform facing an open arm and allowed to explore freely for 10 min. Entries were defined as all four limbs entering an arm. Anxiety-like behaviour was assessed by the percentage of time spent in open arms and the number of open arm entries. Total distance travelled was recorded to control for locomotor activity.

### Open field test (OFT)

The open field test was performed in a 50 × 50 × 50 cm^3^ arena. Mice were placed in the centre and allowed to explore for 20 min. Anxiety-like behaviour was evaluated by time spent in the centre zone and the number of centre entries. Total locomotion distance was used as a control for general activity.

### Tail suspension test (TST)

Mice were suspended by the tail using adhesive tape positioned ∼1-2 cm from the tip and recorded for 6 min. Immobility was defined as the absence of active escape-related movements. Total immobility time was quantified as an index of depressive-like behaviour. Scoring was performed using automated software and confirmed by blinded observers.

### Cell culture and transfection

HGC27 cells were cultured in the logarithmic growth phase and passaged at a 1:3 -1:5 ratio. For siRNA transfection, cells were seeded 24 h prior to achieve approximately 50 - 70% confluency. Three siRNAs targeting NPY (siNPY-1: 5’-CGCUGCGACACUACAUCAATT-UUGAUGUAGUGUCGCAGCGTT-3′; siNPY-2: 5′-GAUGAGAGAAAGCACAGAATT-UUCUGUGCUUUCUCUCAUCTT-3′; siNPY-3: 5′-CGGCUUGAAGACCCUGCAATT- UUGCAGGGUCUUCAAGCCGTT -3′) or NPY1R (siNPY1R-1: 5′-GAGGCGAUGUGUAAGUUGATT- UCAACUUACACAUCGCCUCTT-3′; siNPY1R-2:5′- GGAACGACAUCAGCUGAUATT-UAUCAGCUGAUGUCGUUCCTT-3′; siNPY1R-3: 5′-CAAGAUAUAUAUACGCCUATT- UAGGCGUAUAUAUAUCUUGTT-3′) were transfected at a final concentration of 20□nM using Lipofectamine 3000 (Thermo, L3000008) following the manufacturer’s protocol. The medium was replaced with fresh complete medium 8□h post-transfection, and cells were collected 24□h later for downstream analyses to identify the one with the highest knockdown efficiency.

### Immunohistochemistry

Paraffin-embedded human sections were first deparaffinised, then incubated with 3% H□O□to quench endogenous peroxidase activity, followed by antigen retrieval using a citric acid buffer. After blocking with 2% BSA in TBST, the sections were incubated with primary antibodies for 2 h at 37°C and subsequently overnight at 4 °C. Following thorough washing, sections were incubated with fluorophore-conjugated secondary antibodies. Detailed information on these gastric cancer patients is provided in **Supplementary Table 1**. The primary antibodies used were as follows: Rabbit anti-NPY1R (Invitrogen, PA5-106849), Mouse anti-NPY (Invitrogen, ABS 028-08-02), Rabbit anti- MAP2 (CST, 4542), Rabbit anti-TH (Oasis Biofarm, OB-PRB100-02), Mouse anti-NPY (Invitrogen, ABS 028-08-02), Mouse anti-ChAT (Invitrogen, PAS-29653).

### Immunofluorescence

Mice were anesthetised via intraperitoneal administration of 20% urethane (10 μL/g), followed by transcranial perfusion with PBS and 4% PFA. Brains were carefully dissected, post-fixed in 4% PFA at 4 °C overnight, and subsequently cryoprotected in 30% sucrose solution. Tissues were then sectioned into 20 μm slices using a cryostat. For immunostaining, brain sections were permeabilised in 0.5% Triton X-100 in PBS (PBST) for 1 h, blocked with 2% BSA in PBST for 1 h, and incubated with primary antibodies at 4 °C overnight. After washing, sections were incubated with appropriate secondary antibodies, counterstained with DAPI, and mounted for imaging. The primary antibodies included Rabbit anti- Cleaved caspase-3 (1:500, CST, 9664), Rabbit anti-NPY1R (1:200, Invitrogen, PA5-106849), Rabbit anti- Ki67 (1:1000, CST, 9126), Rabbit anti- GABA (1:500, Sigma, A2052), Rabbit anti- c-Fos (1:500, Sigma, ABE457), Rat anti- NeuN (1:500, Oasis Biofarm, OB-PRT013-01). TUNEL staining was carried out using the In Situ Cell Death Detection Kit, TMR red (Roche, 12156792910) in accordance with the manufacturer’s instructions, and subsequent immunostaining procedures were performed after TUNEL labelling. Image acquisition was performed using a VS200 slide scanner or a SpinSR spinning disk confocal system (Olympus).

### ELISA

Protein levels in plasma or tissue samples were quantified using sandwich ELISA kits according to the manufacturer’s instructions. Samples and standards were run in duplicate or triplicate. Absorbance was measured at 450 nm using a microplate reader. Concentrations were calculated based on standard curves. All samples were assayed within the linear detection range. The following ELISA kits were used: Human anti- NPY (Elabscience, E-EL-H1893c), Mouse anti- NPY (Elabscience, E-EL-H1893c), Human anti- cortisol (Elabscience, E-OSEL-H0006), Mouse anti-cortisol (Fine Test, EM1721), NA/NE (Noradrenaline/Norepinephrine) (Elabscience, E-EL-0047), Acetylcholine (Elabscience, E-EL-0081).

### RNA sequencing and bioinformatics analysis

Total RNA was extracted and assessed for integrity (RIN ≥ 7.0). Libraries were constructed using poly(A) enrichment and sequenced on an Illumina platform (paired-end, 150 bp). Raw reads were quality-filtered and aligned to the reference genome using HISAT2 or STAR. Gene expression levels were quantified and normalised as TPM or FPKM. Differential expression analysis was performed using DESeq2 or edgeR with thresholds: log□fold change ≥ 1 and FDR < 0.05. Functional enrichment analysis (GO and KEGG) was conducted using standard statistical frameworks. Principal component analysis and clustering were used to assess sample variability.

### Western blotting

Proteins were extracted using RIPA buffer (Epizyme, PC101) supplemented with protease and phosphatase inhibitors. Equal amounts of protein were separated by SDS-PAGE and transferred onto PVDF membranes. Membranes were blocked with 5% BSA and incubated with primary antibodies overnight at 4 °C, followed by HRP-conjugated secondary antibodies. The primary antibodies included Rabbit anti-NPY1R antibody (1:1000, Huaan Biotechnology, ER64221). Rabbit anti-p44/42 MAPK (Erk1/2) (1:1000, CST, 4695), Rabbit anti-total Erk1/2 (42/44) (1:1000, CST, 4695), HRP-Conjugated GAPDH (1:5000, Proteintech, HRP-60004). Signals were detected with SuperSignal West Atto Ultimate Sensitivity Substrate (Thermo Fisher Scientific, A38555) by ChemiDoc MP Imaging System (Bio-Rad, USA) and quantified using Fiji. NPY1R protein levels were normalised to GAPDH, and p44/42 MAPK (Erk1/2) protein levels were normalised to total Erk1/2 (42/44).

### Quantitative real-time PCR (qPCR)

Total RNA was isolated using RNAzol reagent (Sigma-Aldrich, R4533), and cDNA was generated using the Evo M-MLV reverse transcription kit (Accurate Biology, AG11706) following the manufacturer’s protocol. qPCR was performed with the SYBR Green Premix Pro Taq HS kit (Accurate Biology, AG11701). Primer sequences are provided in Supplementary Table 1. Gene expression levels were quantified using the 2^-ΔΔCt method, with GAPDH serving as the internal reference. Data are expressed as fold changes relative to the control group. Primers for qPCR are as follows: NPY1R (Homo sapiens): 5′-GCAGGAGAAATACCAGCGGA-TCCCTTGAACTGAACAATCCTC-3′; NPY (Homo sapiens): 5′-CGCTGCGACACTACATCAAC- CTCTGGGCTGGATCGTTTTCC-3′; GAPDH (Homo sapiens): 5′-TGTGGGCATCAATGGATTTGG- -ACACCATGTATTCCGGGTCAAT-3′.

### Bioinformatic Analysis and Prognostic Evaluation

Data from The Cancer Genome Atlas were used to analyse stomach adenocarcinoma (STAD) samples, focusing on the expression profiles of NPY and its receptor subtypes (NPY1R, NPY2R, NPY4R, NPY4R2, NPY5R, and NPY6R) in gastric cancer tissues. Correlation analysis was performed between gene expression levels and tumour grade. Survival curves were generated using the Kaplan-Meier method, and the impact of NPY1R expression on overall survival (OS) was evaluated using the Log-rank test.

### Statistical analysis

Data are presented as mean ± SEM. Statistical analyses were performed using GraphPad Prism. Two-group comparisons were analysed using a two-tailed Student’s t-test. Multiple comparisons were performed using one-way ANOVA or two-way ANOVA followed by Tukey’s post hoc tests. No statistical methods were used to predetermine sample size. Sample sizes were chosen based on previous studies and pilot experiments. All *in vitro* experiments were independently repeated at least three times. A *p* < 0.05 was considered statistically significant.

## Data availability

All other data supporting the findings of this study are available from the corresponding author on reasonable request.

## Acknowledgements

This work was supported by Shenzhen Fundamental Research Program (RCJC20231211090018040 and ZDSYS20220606100801003 to C.Y.; JCYJ20240813150421029 to X.H.), National Natural Science Foundation of China (W2511095 and 32271034 to J.N.).

## Author contributions

C.Y. and J.N. conceptualized the study, designed the experiments, and composed the paper. X.H. conducted most of the experiments. Y.Z. conducted most of the animal experiments. Q.W. assisted with the bioinformatics analysis. C.W. and S.R. conducted the electrophysiological recording. X.N., Y.H., and C.Z. contributed to human samples collection. J.W assisted with the animal experiments. R.F., L.L., B.O., H.C. and A.V. contributed to the interpretation of the results. R.F., L.L., B.O., H.C. and A.V. edited and reviewed the manuscript.

---

**Extended Data Fig. 1.**
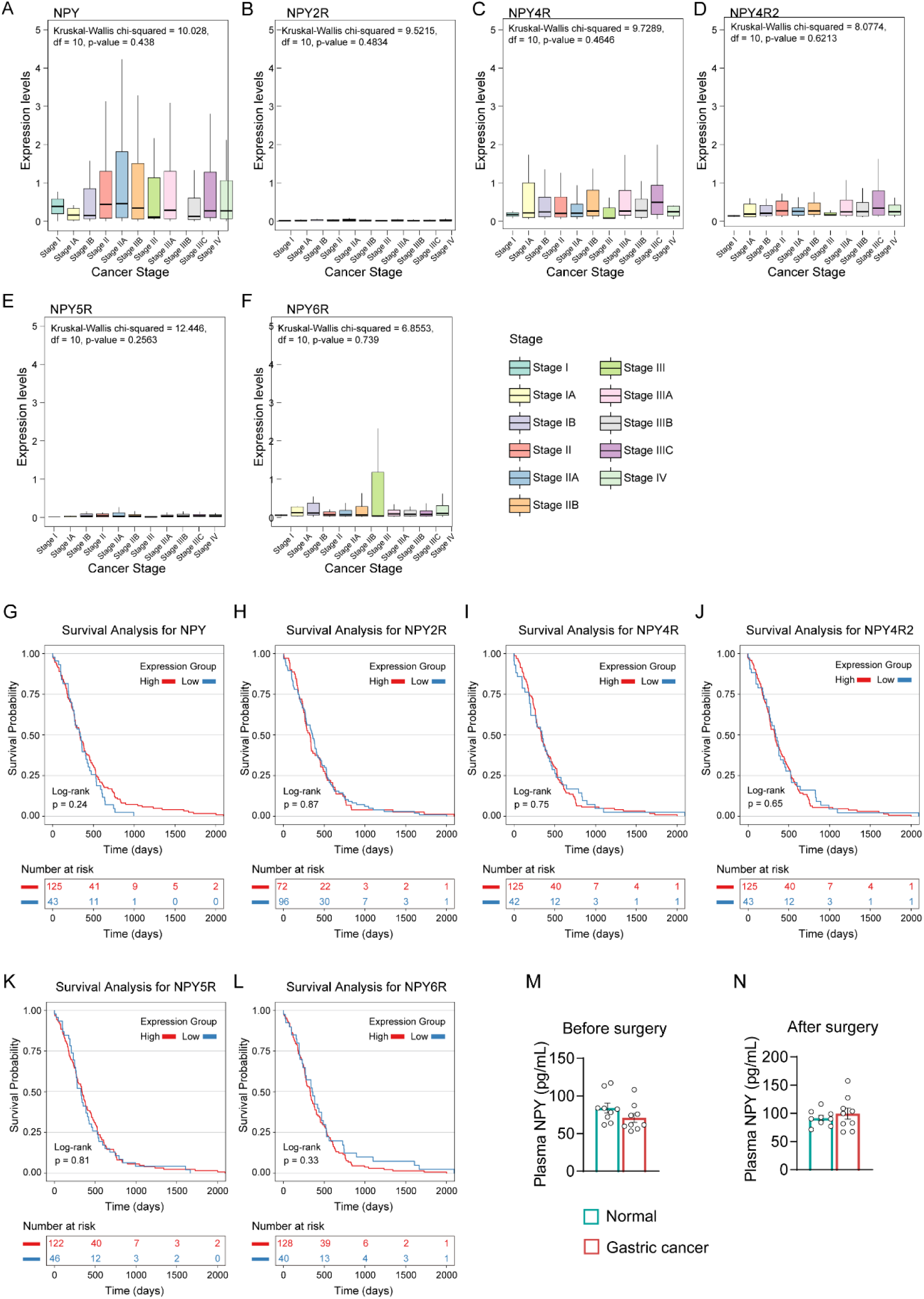
Expression of NPY and its receptor subtypes in gastric cancer tissues at different TNM clinical stages and their association with the survival rate of gastric cancer patients. (A-F) Expression levels of NPY and its receptor subtypes across different TNM clinical stages in the STAD cohort. (G-L) Kaplan-Meier survival curves from the STAD cohort showing the correlation between NPY signalling-related molecules and overall survival in gastric cancer patients. (M-N) Comparison of plasma NPY levels between gastric cancer patients and healthy controls before and after surgery.

**Extended Data Fig. 2.**
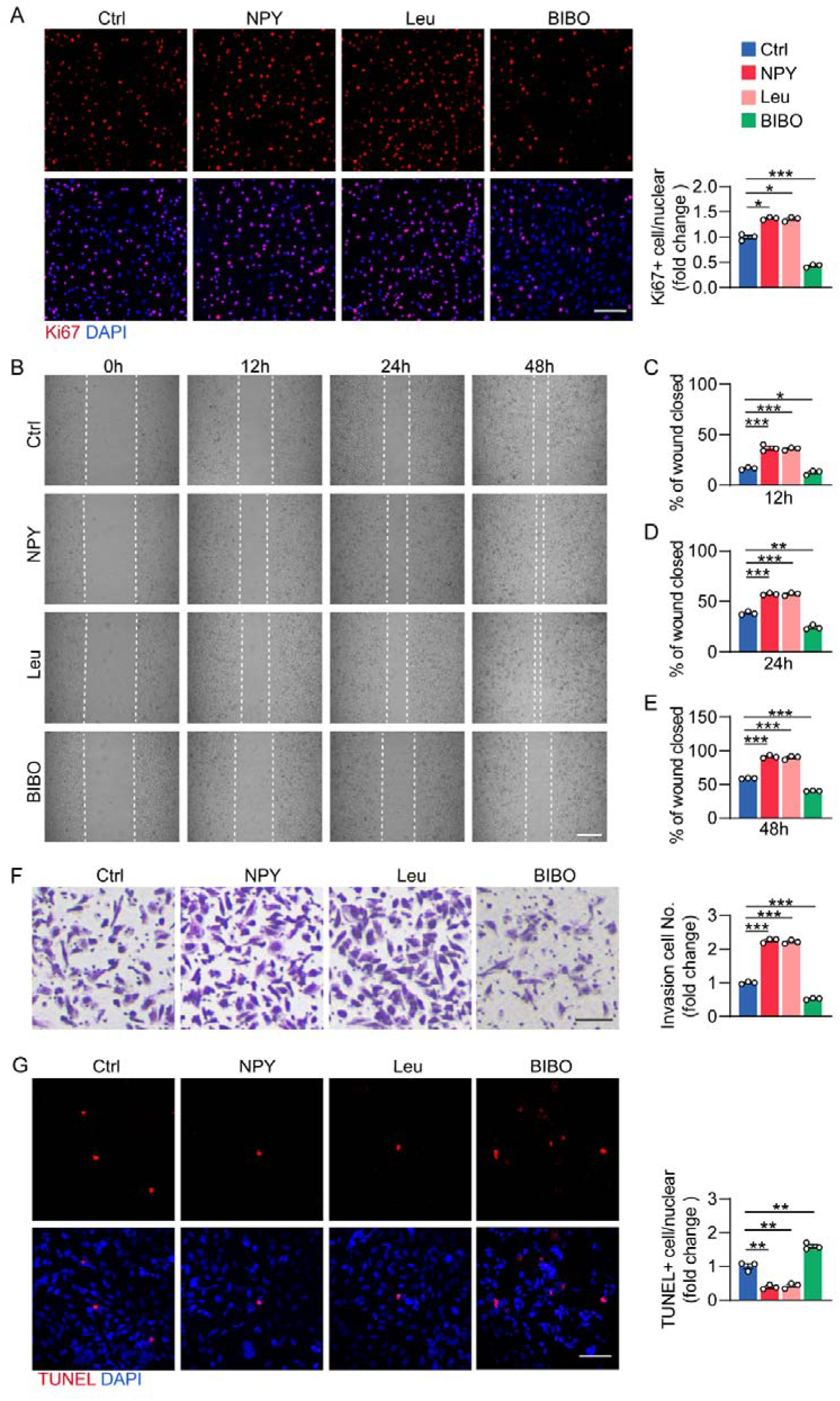
*In vitro* functional experiments on HGC27 cells validating the direct effects of NPY/NPY1R signalling on gastric cancer cell behaviour. HGC27 gastric cancer cells were treated with NPY, NPY1R agonist [Leu31, Pro34]-Neuropeptide Y (Leu), or the NPY1R antagonist BIBO. (A) Representative immunofluorescence images of KI67-positive cells. Scale bar: 100 μm. Right: quantification of KI67-positive cells. (B) Wound healing assays were performed to evaluate HGC27 cell migration at 12 h, 24 h, and 48 h. Scale bar: 500 μm. Right: quantification of wound closed ratio at (C) 12 h, (D) 24 h, and (E) 48 h. (F) Representative images of invasion assays to test the invasive capacity of HGC27 cells. Scale bar: 50 μm. Right: quantification of invasive cell number. (G) Representative images of TUNEL staining. Scale bar: 50 μm. Right: quantification of TUNEL-positive cells. Data are presented as mean ± SEM, and were analysed by one-way ANOVA with Tukey’s post hoc test, n = 3 independent experiments, \**p* < 0.05, \*\**p* < 0.01, \*\*\**p* < 0.001.

**Extended Data Fig. 3.**
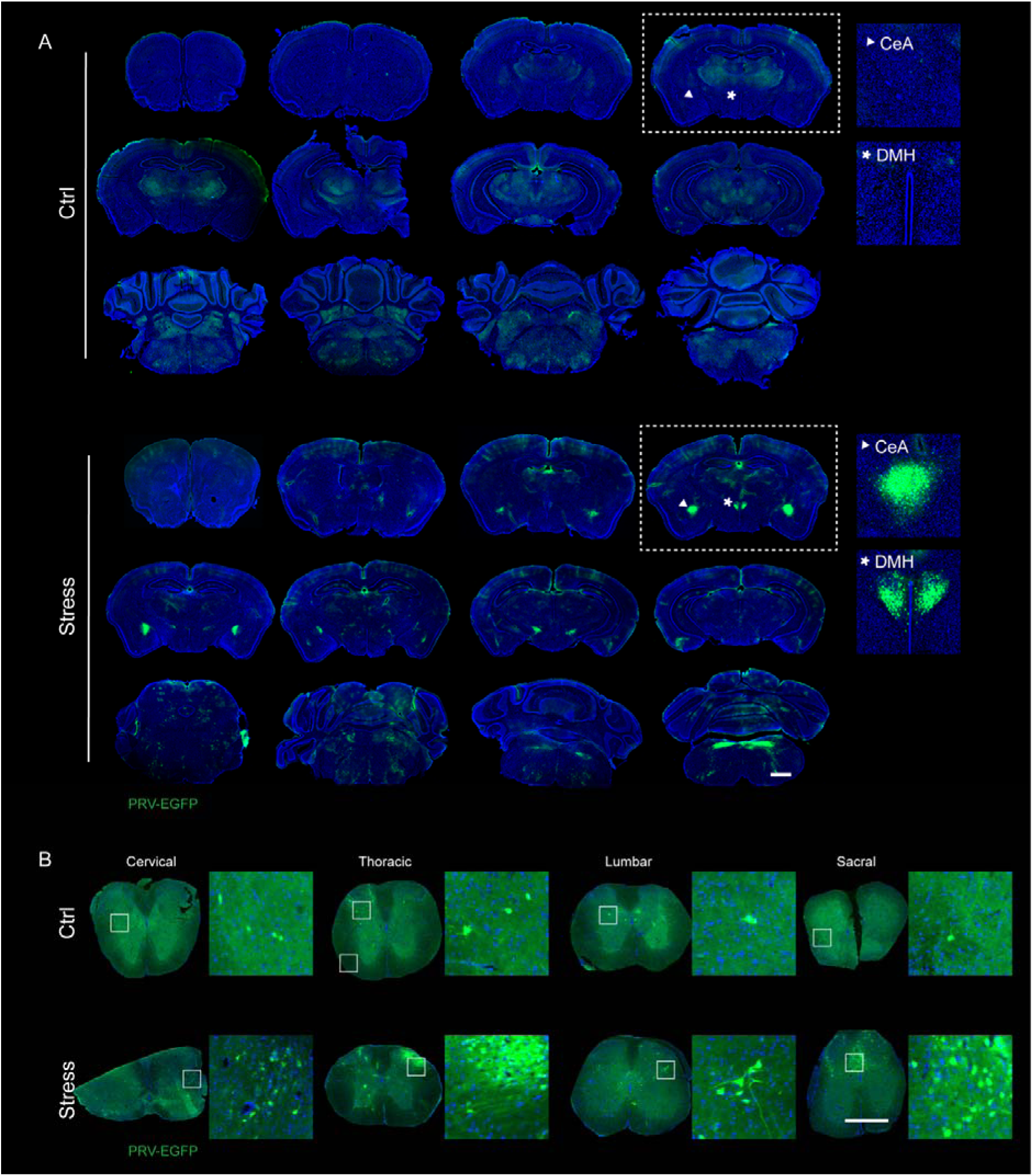
Retrograde trans-multisynaptic viral tracing reveals the projection pathways of tumour-associated central nervous system under chronic stress. (A, B) Representative images of PRV-EGFP signals in the brain sections and spinal cord from ctrl and chronic stress tumour-bearing mice. PRV-EGFP originating from the tumour site was detected transmitting stepwise along the dorsal horn of the sacral, lumbar, thoracic, and cervical spinal cord, demonstrating polysynaptic connectivity between the tumour and the central nervous system. Scale bars: 1 mm.

**Extended Data Fig. 4.**
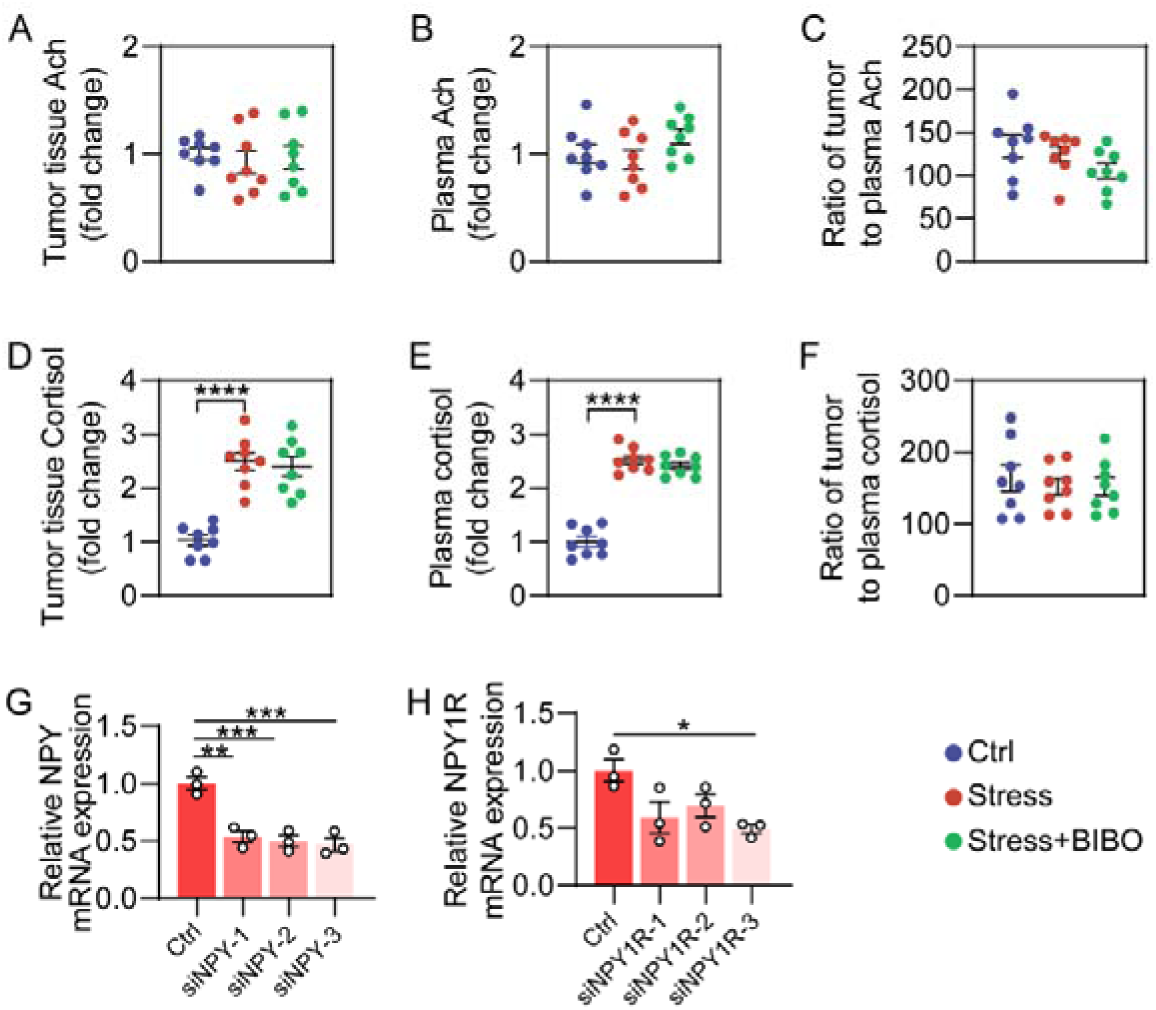
Effects of chronic stress on Ach and cortisol levels in the tumour microenvironment. (A, B) Acetylcholine (Ach) levels in tumour tissues and plasma of ctrl, stress, and stress+BIBO tumour-bearing mice. n = 8. (C) The ratio of Ach in Tumour tissue/plasma in ctrl, stress, and stress+BIBO tumour-bearing mice. n = 8. (D, E) Cortisol levels in tumour tissues and plasma of ctrl, stress, and stress+BIBO tumour-bearing mice. n = 8 mice. (F) The ratio of Cortisol levels in Tumour tissue/plasma in ctrl, stress, and stress+BIBO tumour-bearing mice. n = 8. (G, H) NPY and NPY1R mRNA levels after transfection with siRNAs. n = 3 independent experiments. Data are presented as mean ± SEM, and were analysed by one-way ANOVA with Tukey’s post hoc test, \**p* < 0.05, \*\**p* < 0.01, \*\*\**p* < 0.001, \*\*\*\**p* < 0.0001.

**Extended Data Fig. 5.**
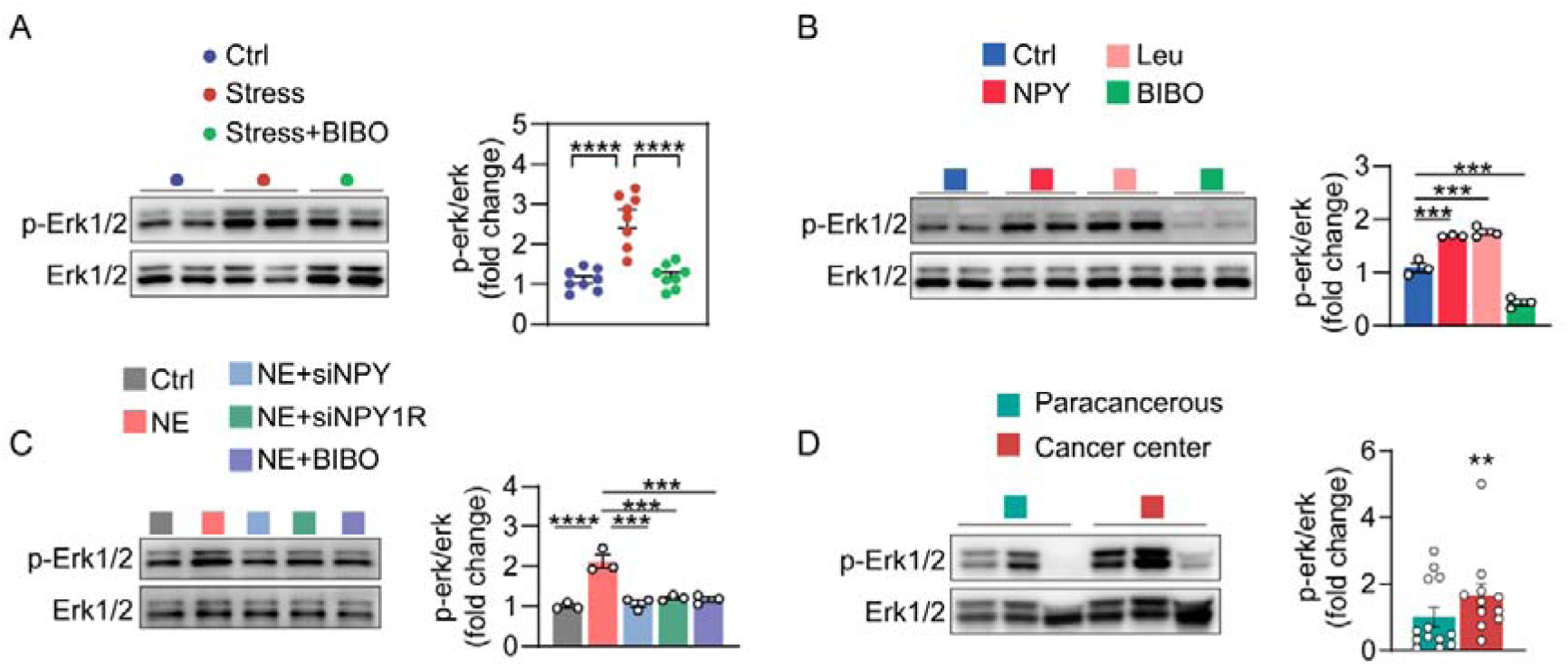
The NPY/NPY1R axis promotes tumour progression through activation of the MAPK signalling pathway. (A) p-Erk1/2 levels in tumour tissues of mice within chronic restraint stress by Western blot. n = 8. (B) Effects of NPY, NPY1R agonist (Leu), or NPY1R inhibitor BIBO on p-Erk1/2 levels in HGC27 cells by Western blot. n = 3 independent experiments. (C) Western blot analysis of p-Erk1/2 levels in NE treated HGC27 cells with or without siRNA/BIBO treatment. n = 3 independent experiments. (D) p-Erk1/2 levels between human gastric cancer tissues and adjacent non-cancerous tissues by Western blot. n = 11-13. Data are presented as mean ± SEM. Data in A-C were analysed by one-way ANOVA with Tukey’s post hoc test and data in D were analysed by two-tailed *t* test, \**p* < 0.05, \*\**p* < 0.01, \*\*\**p* < 0.001, \*\*\*\**p* < 0.0001.

**Extended Data Fig. 6.**
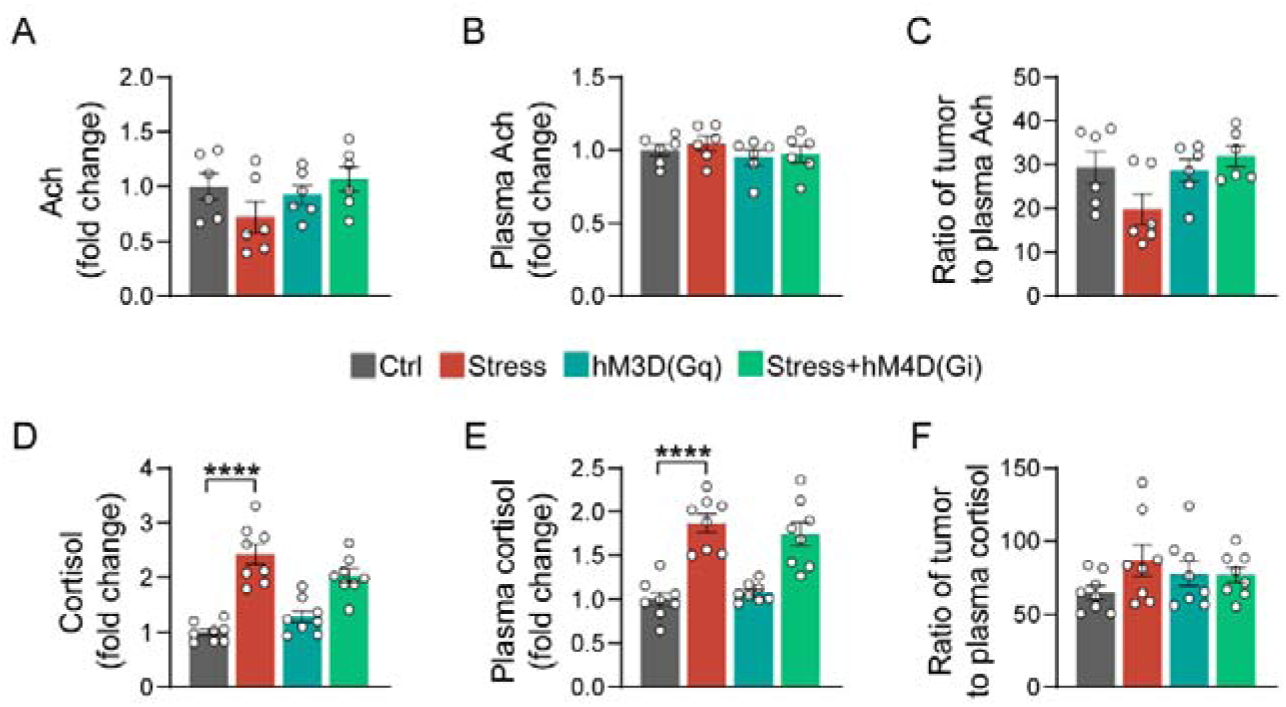
Effects of manipulating CeA activity on acetylcholine and cortisol levels in tumour tissue and plasma. (A, B) Acetylcholine (Ach) levels in tumour tissues and plasma from Ctrl, Stress, hM3D(Gq), and Stress+hM4D(Gi) tumour-bearing mice. (C) Ratio of Ach levels in tumour/plasma. (D, E) Cortisol levels in tumour tissues and plasma from Ctrl, Stress, hM3D(Gq), and Stress+hM4D(Gi) tumour-bearing mice. Data were analysed by one-way ANOVA with Tukey’s post hoc test. (F) Ratio of tumour tissue to plasma cortisol levels in indicated groups. Data are presented as mean ± SEM (n = 6-8), and were analysed by one-way ANOVA with Tukey’s post hoc test, \*\*\**p* < 0.001, \*\*\*\**p* < 0.0001.

